# Rdh10-mediated Retinoic Acid Signaling Regulates the Neural Crest Cell Microenvironment During ENS Formation

**DOI:** 10.1101/2025.01.23.634504

**Authors:** Naomi E. Butler Tjaden, Stephen R. Shannon, Christopher W. Seidel, Melissa Childers, Kazushi Aoto, Lisa L. Sandell, Paul A. Trainor

## Abstract

The enteric nervous system (ENS) is formed from vagal neural crest cells (NCC), which generate most of the neurons and glia that regulate gastrointestinal function. Defects in the migration or differentiation of NCC in the gut can result in gastrointestinal disorders such as Hirschsprung disease (HSCR). Although mutations in many genes have been associated with the etiology of HSCR, a significant proportion of affected individuals have an undetermined genetic diagnosis. Therefore, it’s important to identify new genes, modifiers and environmental factors that regulate ENS development and disease. Rdh10 catalyzes the first oxidative step in the metabolism of vitamin A to its active metabolite, RA, and is therefore a central regulator of vitamin A metabolism and retinoic acid (RA) synthesis during embryogenesis. We discovered that *retinol dehydrogenase 10* (*Rdh10*) loss-of-function mouse embryos exhibit intestinal aganglionosis, characteristic of HSCR. Vagal NCC form and migrate in *Rdh10* mutant embryos but fail to invade the foregut. *Rdh10* is highly expressed in the mesenchyme surrounding the entrance to the foregut and is essential between E7.5-E9.5 for NCC invasion into the gut. Comparative RNA-sequencing revealed downregulation of the *Ret-Gdnf-Gfrα1* gene signaling network in *Rdh10* mutants, which is critical for vagal NCC chemotaxis. Furthermore, the composition of the extracellular matrix through which NCC migrate is also altered, in part by increased collagen deposition. Collectively this restricts NCC entry into the gut, demonstrating that *Rdh10*-mediated vitamin A metabolism and RA signaling pleiotropically regulates the NCC microenvironment during ENS formation and in the pathogenesis of intestinal aganglionosis.

## Introduction

The gastrointestinal tract is an endoderm derived organ system that functions to process foods and liquids via mechanical and chemical digestion, while also playing a major role in immune system recognition of, and in response to, introduced pathogens(1–6). Most functions of the gastrointestinal tract are governed by the enteric nervous system (ENS) which is derived from vagal neural crest cells (NCC). Beginning around embryonic day (E) 8.75-9.0 of mouse embryogenesis, these multipotent progenitor cells delaminate from the neuroepithelium at the axial level of somites 1-7, migrate ventrally through the adjacent mesoderm, and enter the foregut mesenchyme by E9.5. Once in the gut, NCC must proliferate, migrate and colonize the full length of the growing gastrointestinal tract, and differentiate into the neurons and glia that comprise the myenteric and submucosal plexi of the ENS. Defects in NCC development can result in the absence of enteric ganglion cells along variable portions of the gastrointestinal tract, disrupting gut function and peristalsis. This type of gastrointestinal disorder is termed Hirschsprung disease (HSCR)(7), and affects 1 in 5000 live births. Patients with HSCR typically present with an inability to pass stool, resulting in abdominal discomfort and distension. Currently, treatment involves surgical resection of the aganglionic bowel(8), but patients with total intestinal aganglionosis require intestinal transplantation. Cell-based therapies for enteric neuropathies such as HSCR have emerged as potential, future treatment options that may one day be used in place of surgical intervention with more favorable patient outcomes (9–14). However, one of the greatest challenges to the development of new preventative therapies for HSCR is its complex etiology, being both multi-genic and variably penetrant. Although mutations in the *RET-GDNF-GFRα1* signaling pathway genes account for 15%-35% of sporadic HSCR and about 50% of familial cases(15), a significant proportion of affected individuals have no known genetic diagnosis. Therefore, additional genes, modifiers and environmental factors must play important roles in the etiology of Hirschsprung disease, so it remains important to identify and understand the signals and mechanisms that regulate vagal neural crest cell development(16, 17).

To discover novel genes that play important roles in neural crest cell development in an unbiased manner, we performed a three generation forward genetic screen in mice via N-ethyl-N-nitrosourea (ENU) induced mutagenesis(18). From this screen we recovered a mouse mutant, termed *trex*, which exhibited craniofacial(19–21) and limb malformations(22). Subsequently, we identified a point mutation in *retinol dehydrogenase 10* (*Rdh10*) underlying the etiology of the *trex* phenotype. Rdh10 catalyzes the first oxidation reaction in the metabolism of vitamin A to its active metabolite, retinoic acid (RA). More specifically, retinol binding protein 4 (RBP4) transports vitamin A (all trans-retinol) from hepatocyte storage into cells via the cell surface receptor Stra6, and therein RDH10 converts all-trans-retinol to all-trans-retinal(19, 23, 24). All-trans-retinal is then oxidized to RA by ALDH1A1, ALDH1A2 and ALDH1A3 (formerly RALDH1, RALDH2, RALDH3) (25–27). RA is an essential signaling molecule in embryonic development, and through its interaction with retinoic acid receptors (RARs) and retinoid receptors (RXRs), which bind as heterodimers to the promoters of developmental genes, RA controls gene transcriptional activity(28–31). Both excess and deficiency of vitamin A and RA can cause developmental malformations and therefore physiological levels must be precisely controlled.

Using an allelic series of *Rdh10* mice, we investigated both spatial and temporal contributions of *Rdh10*-mediated vitamin A metabolism and RA signaling to vagal NCC development and ENS formation during gastrointestinal development. We discovered that *Rdh10* is required in a paracrine fashion between E7.5-9.5 for NCC migration, entry and colonization of the gut, and that loss-of-function results in intestinal aganglionosis. Comparative RNA-sequencing revealed that *Rdh10*-mediated RA signaling positively regulates the *Ret-Gdnf-Gfrα1* gene network, which is critical for chemotactic entry of vagal NCC into the foregut and their subsequent migration and differentiation. RA signaling also regulates extracellular matrix composition (ECM), and in *Rdh10* mutants, we observed major changes such as increased collagen deposition, possibly in association with increased stiffness. Together with alterations in focal cell adhesion, this may collectively impede NCC migration and prevent their entry into the gut, demonstrating that *Rdh10*-mediated vitamin A metabolism and RA signaling pleiotropically regulate the NCC microenvironment during ENS formation. *Rdh10* loss-of-function and vitamin A deficiency therefore represent genetic and non-genetic risk factors respectively, in the etiology, penetrance and severity of intestinal aganglionosis and possibly Hirschsprung disease.

## Results

### *Rdh10^trex/trex^* embryos exhibit intestinal aganglionosis

*Rdh10^trex/trex^* mutant embryos were recovered in an ENU mutagenesis screen designed to identify novel genes important for the development of neural crest cells and their derivatives(18). In characterizing the craniofacial and limb malformations (18, 19, 21, 32), we performed β-tubulin III (TUJ1) immunostaining to assay for neuronal differentiation (Figure 1A-D). E10.5 wild-type embryos exhibit extensive neuronal differentiation in the forebrain and midbrain with well-developed cranial and dorsal root ganglion peripheral nerve projections (Figure 1A). In contrast, *Rdh10^trex/trex^* embryos presented with hypoplastic forebrain and midbrain neural networks, abnormal cranial nerve patterning, and an absence of enteric neurons in the developing GI tract (Figure 1C). The absence of mature enteric neuronal differentiation is clearly apparent in isolated intestines from E11.5 *Rdh10^trex/trex^* embryos compared to controls (Figure 1B,D). This intestinal aganglionosis phenotype mirrors severe Hirschsprung Disease (HSCR) in humans, rendering *Rdh10^trex/trex^* embryos a potential new genetic model of this devastating condition.

**Figure 1.**
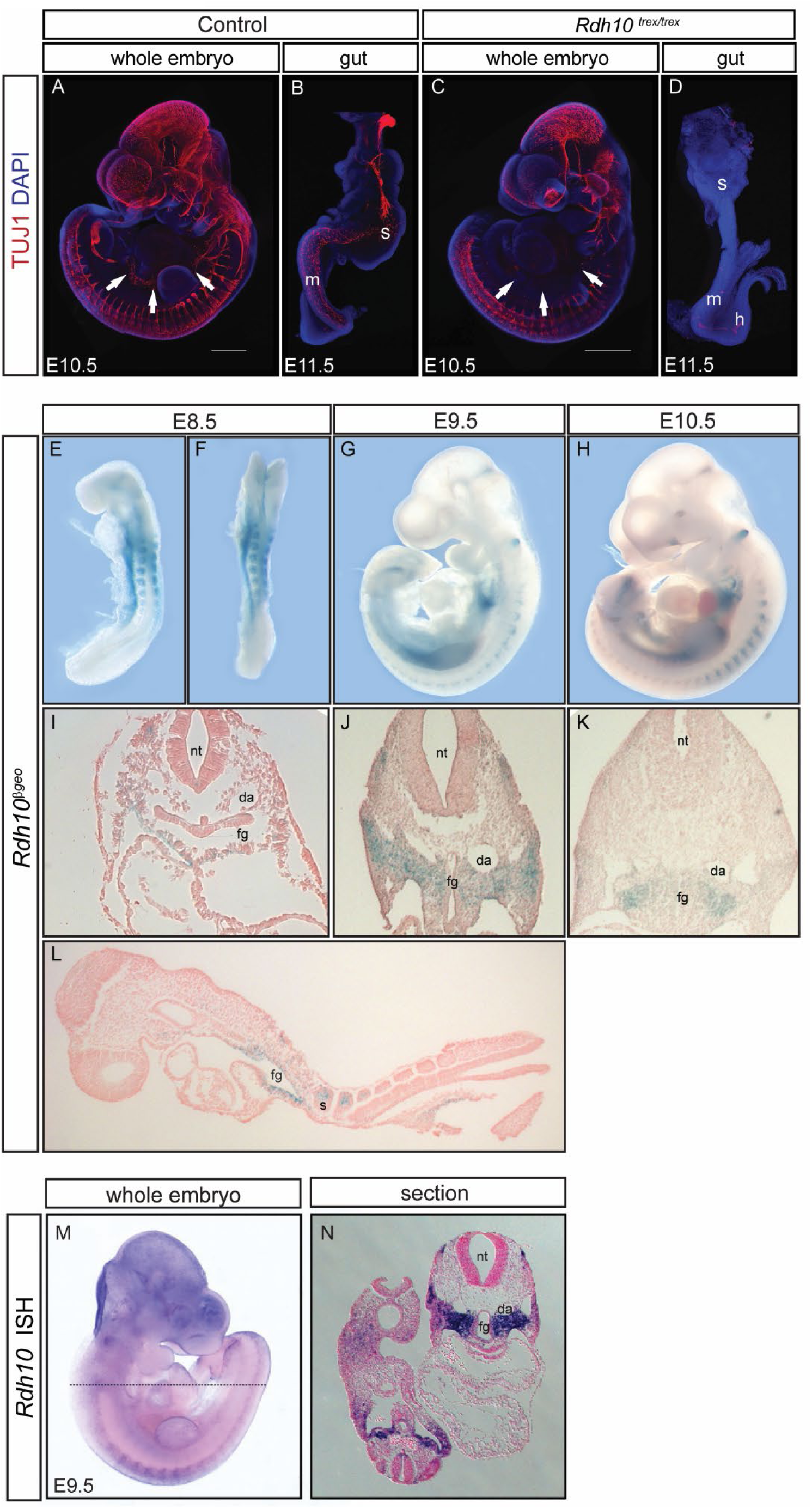
*Rdh10^trex^*is a model for Hirschsprung Disease. TUJ1 (red) and DAPI (blue) immunostaining in E10.5 whole embryos (A,C) and E11.5 whole guts (B,D). (A,C) Mature neurons are labeled with TUJ throughout the head and trunk of *Rdh10^trex/trex^* mutant and control littermates. White arrows (A) denote the developing enteric nervous system. Arrows in the mutant embryo (C) indicate intestinal aganglionosis with the absence of enteric neurons. (B) Mature neurons in the E11.5 control gut are present in the stomach and midgut extending to the cecum. (D) The mutant gut shows complete neuronal agenesis with a lack of discernable staining in the stomach, midgut and hindgut. (E-L) Spatiotemporal expression of *Rdh10* during normal embryogenesis between E8.5-10.5 shown by LacZ staining of the *Rdh10^βgeo^* mouse line. At E8.5, lateral (E) and ventral (F) views of the whole embryo as well as transverse (I) and sagittal (L) sections show *Rdh10* expression in the anterior somites, lateral mesoderm and in the mesenchyme surrounding the primitive gut. At E9.5, views of the lateral whole embryo (G) and transverse section (J) show *Rdh10* expression in the somites, ventral mesoderm, and anterior foregut diverticulum mesenchyme. At E10.5, views of the lateral embryo (H) and transverse section (K) show *Rdh10* expression in the somites, ventral mesoderm, and anterior foregut. *Rdh10* mRNA expression by in situ hybridization is seen in a lateral view of the whole embryo (M) and transverse section (N) within the somites, ventral mesoderm and anterior foregut mesenchyme. Scale bars are 500μm. (s) stomach; (m) midgut; (h) hindgut; (nt) neural tube; (da) dorsal aorta; (fg) foregut; (s) somite.

As a first step in understanding the mechanistic origins of the gastrointestinal defects in *Rdh10^trex/trex^* embryos, we characterized the spatiotemporal activity of *Rdh10* during normal embryogenesis (Figure 1E-N). We previously reported that *Rdh10* expression is first detected at E8.0, in the neural groove of the prospective hindbrain and in the lateral mesoderm(19). Using *Rdh10*^βgeo/+^ mice where LacZ activity serves as a proxy for *Rdh10* expression, we observed LacZ staining in the anterior somites and adjacent lateral mesoderm as well as in the mesenchyme surrounding the primitive gut at E8.5 (Figure 1E,F,I,L). The axial domain of staining corresponds to the vagal region from which enteric NCC are derived. The spatiotemporal expression of *Rdh10* is developmentally dynamic as is evidenced by the pattern of LacZ staining in E9.5-10.5 embryos. *Rdh10* is expressed in posterior somites in E9.5 embryos and continues to be expressed in the ventral mesoderm as well as the anterior foregut diverticulum mesenchyme (Figure 1G,J). A similar pattern of *Rdh10* expression is observed via *in situ* hybridization (Figure 1M,N). In E10.5 *Rdh10*^βgeo/+^ embryos, LacZ activity persists in the somites and ventral mesoderm, but has decreased considerably in the mesenchyme at the entrance to the foregut (Figure 1H,K). Interestingly, the spatiotemporal expression of *Rdh10* encompasses the ventral migration route vagal neural crest cells traverse from the neural tube to the foregut, implying it could play a critical role in NCC migration or entry into the foregut and their subsequent formation of the ENS during gastrointestinal development.

### Vagal NCCs migrate ventrally but do not invade the foregut of *Rdh10^trex^*^/^*^trex^* embryos

To determine whether perturbation of NCC migration and/or differentiation could account for intestinal aganglionosis in *Rdh10^trex/trex^* embryos, we examined the activity of *Sox10* (Supplemental Figure 1A-D’), which is expressed by migrating NCC and is a master regulator of neuroglial differentiation. In E9.5 control embryos, *Sox10* staining indicates NCC form and migrate ventrally into the anterior region of the foregut (Supplemental Figure 1A,A’). This differs considerably from *Rdh10^trex/trex^* mutant embryos in which *Sox10^+^* NCC remain in closer proximity to the neural tube, with diminished ventral migration that culminates in an absence of NCC within the foregut region (Supplemental Figure 1C,C’). By E10.5, a stream of *Sox10^+^* cells extending caudally into the midgut is evident in the GI tract of control embryos (Supplemental Figure 1B,B’). In contrast, *Rdh10^trex/trex^* mutant embryos continue to exhibit a complete absence of vagal NCC in the foregut (Supplemental Figure 1D,D’). Although *Sox10^+^* NCC are formed in *Rdh10^trex/trex^*embryos, their ventral migration is disrupted between E9.0-E10.5, and they fail to invade the foregut or colonize more caudal regions of the GI tract.

To validate perturbation of NCC migration as the cellular mechanism underpinning the pathogenesis of intestinal aganglionosis in *Rdh10^trex/trex^* embryos, we labeled migrating NCC and their neuronal derivatives *in vivo* using the *Mef2c-F10N-LacZ* reporter mouse(19) (Figure 2A-L). LacZ staining of control embryos reveals that NCCs migrate ventrally and begin to invade the foregut mesenchyme by E9.5 (Figure 2A-C). In contrast, although neural crest cells migrate ventrally in *Rdh10^trex/trex^* mutant embryos, they remain distant from the foregut (Figure 2D-F). By E10.5, the differences in neural crest cell migration and invasion of the gut mesenchyme are profound. In control embryos, vagal NCC have clearly migrated into the foregut with colonization extending to the midgut (Figure 2G-I). However, in *Rdh10^trex/trex^* embryos, no NCC appear to have invaded the foregut mesenchyme and very few LacZ^+^ NCC reach the vicinity of the foregut (Figure 2J,K). This abnormal pattern of NCC migration and failure to invade the foregut are evident in embryo sections (Figure 2L). These data suggest that vagal NCC form and migrate ventrally through the adjacent mesoderm in *Rdh10^trex/trex^* embryos but fail to invade the foregut mesenchyme to colonize the gastrointestinal tract.

**Figure 2.**
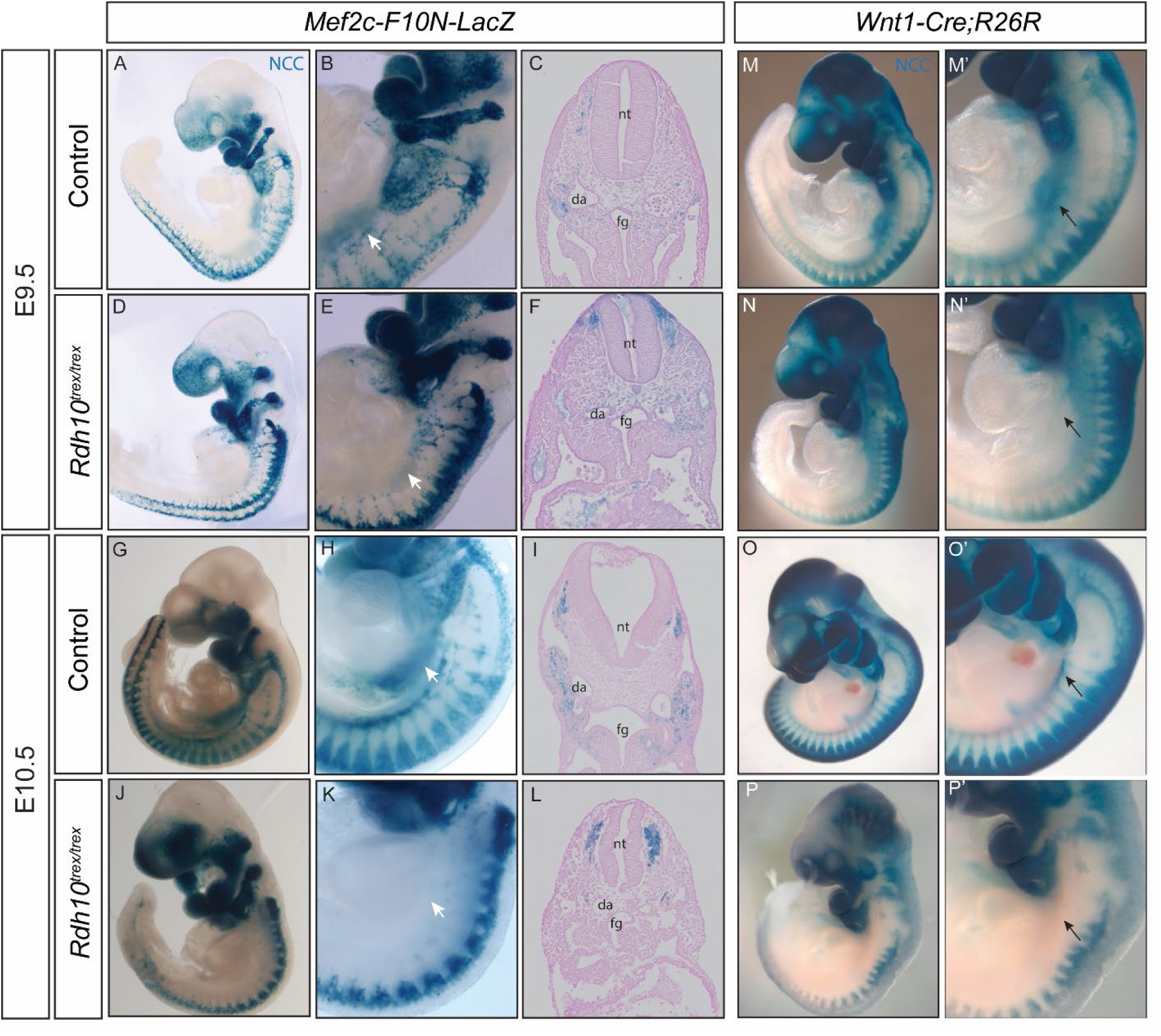
Vagal neural crest cell migration is disrupted in *Rdh10^trex/trex^* mutant embryos. (A-L) LacZ staining of the *F10N* mouse line bred into the *Rdh10^trex/trex^* background label migrating NCC in blue indicated by the white arrowheads (B,E,H,K). At E9.5, lateral view of control whole embryos (A,B) and transverse section (C) reveal staining of a vagal NCC migrating ventrally into the foregut mesenchyme. At E10.5, lateral view of control whole embryos (G,H) and transverse section (I) similarly show NCC migration into the foregut mesenchyme and midgut region. In contrast, at E9.5 *Rdh10^trex/trex^* embryos (D,E) reveal initial vagal NCC migration, and transverse section (F) shows absent neural crest cell staining in the foregut mesenchyme. Similarly, at E10.5, *Rdh10^trex/trex^* mutant embryos (J,K) reveal ventrally migrating NCCs, but transverse section (L) indicate a lack of NCC in the foregut mesenchyme. (M-P’) LacZ staining of the *Wnt1-* cre*;R26R* mouse line bred into the background of *Rdh10^trex/trex^* label pre-migratory NCC and their descendants in blue. (M’-P’) Black arrows indicate the migrating vagal NCC stream in control and *Rdh10^trex/trex^* whole embryos. At E9.5, lateral view of control whole embryo (M,M’) reveals expected vagal NCC staining extending into the foregut. At E10.5, a lateral view of control whole embryo (O,O’) similarly shows staining of the vagal NCC stream extending from the foregut into the midgut region. In contrast, whole *Rdh10^trex/trex^* embryos lack a vagal NCC stream entirely at both E9.5 (N,N’) and E10.5 (P,P’). (nt) neural tube; (da) dorsal aorta; (fg) foregut.

Although *Sox10* and *Mef2c-F10N-LacZ* label migrating NCC and their neuronal derivatives(19), it was important to perform NCC lineage tracing in control and mutant *Rdh10^trex/trex^*embryos to examine the possibility of a switch in cell fate. *Wnt1-Cre;R26R* was bred into *Rdh10*^trex/trex^ embryos to label dorsal neuroepithelial cells which encompass premigratory NCC and their descendants (Figure 2M-P’). In E9.5 control embryos, LacZ labeled vagal NCC migrate ventrally and enter the foregut (Figure 2 M,M’). By E10.5, LacZ labelled NCC form a clear caudally migrating stream in the GI tract extending from the foregut towards the midgut (black arrow in Figure 2O,O’). In contrast, although NCC migrate ventrally in E9.5 *Rdh10^trex/trex^* embryos, the extent of migration is considerably diminished compared to controls and NCC do not enter the foregut region (Figure 2N,N’). By E10.5, the confirmed absence of a caudally migrating stream of LacZ labelled cells in the gastrointestinal tract is indicative of the failure of NCC colonize the foregut in *Rdh10^trex/trex^* embryos (Figure 2P,P’). *Sox10* expression in combination with *Mef2c-F10N-LacZ* and *Wnt1-Cre;R26R* labeling collectively suggests that intestinal aganglionosis in *Rdh10^trex/trex^* embryos is due to the failure of NCC to migrate into the foregut.

To assess for NCC viability we performed immunostaining with Caspase-3 to label apoptotic cells in *Rdh10*^trex/trex^ and control embryos (Supplemental Figure 2A-F’). Although we observed a small amount of cell death in the mesenchyme in close proximity to the foregut in *Rdh10^trex/trex^*embryos compared to controls (Supplemental Figure 2C-D’), it did not coincide with Ap2α+ labeled neural crest cells (Supplemental Figure 2E-F’). Interestingly, *Rdh10* is expressed in the mesenchyme adjacent to the foregut in E9.5 embryos (Figure 1M,N) correlating with the path NCC traverse to enter the foregut. These findings support altered migration as the basis for the absence of vagal NCC in the foregut of *Rdh10^trex/trex^* and implies that *Rdh10*-mediated RA signaling plays a paracrine or extrinsic role in the microenvironment of vagal NCC, facilitating their invasion and colonization of the foregut.

### *Rdh10^trex/trex^* mice display defects in retinoic acid signaling

We previously reported that Rdh10 functions as a nodal point in vitamin A metabolism and the synthesis of RA during early embryogenesis (19, 20, 32). The *trex* mutation renders the enzymatic dehydrogenase/reductase activity of *Rdh10* null and incapable of oxidizing vitamin A (retinol) to retinal(19). Therefore, we hypothesized that the perturbation of neural crest cell migration and the pathogenesis of intestinal aganglionosis in *Rdh10^trex/trex^* embryos was associated with diminished RA signaling. *RARE-LacZ* was then bred into *Rdh10^trex/trex^* embryos to visualize RA signaling via LacZ staining (Figure 3A-D)(19). In E8.5 wild-type embryos RA signaling is broadly active in the neuroepithelium and adjacent ventral mesenchyme from the axial level of the hindbrain extending posteriorly nearly all the way to the tail(19).. In E8.5 wild-type embryos RA signaling is broadly active in the neuroepithelium and adjacent ventral mesenchyme from the axial level of the hindbrain extending posteriorly nearly all the way to the tail(33). By E9.5 RA signaling has become active in the forebrain, while the previous anterior limit of *RARE-LacZ* activity has receded to the hindbrain/spinal cord junction with signaling now extending into the tail (Figure 3A).

**Figure 3.**
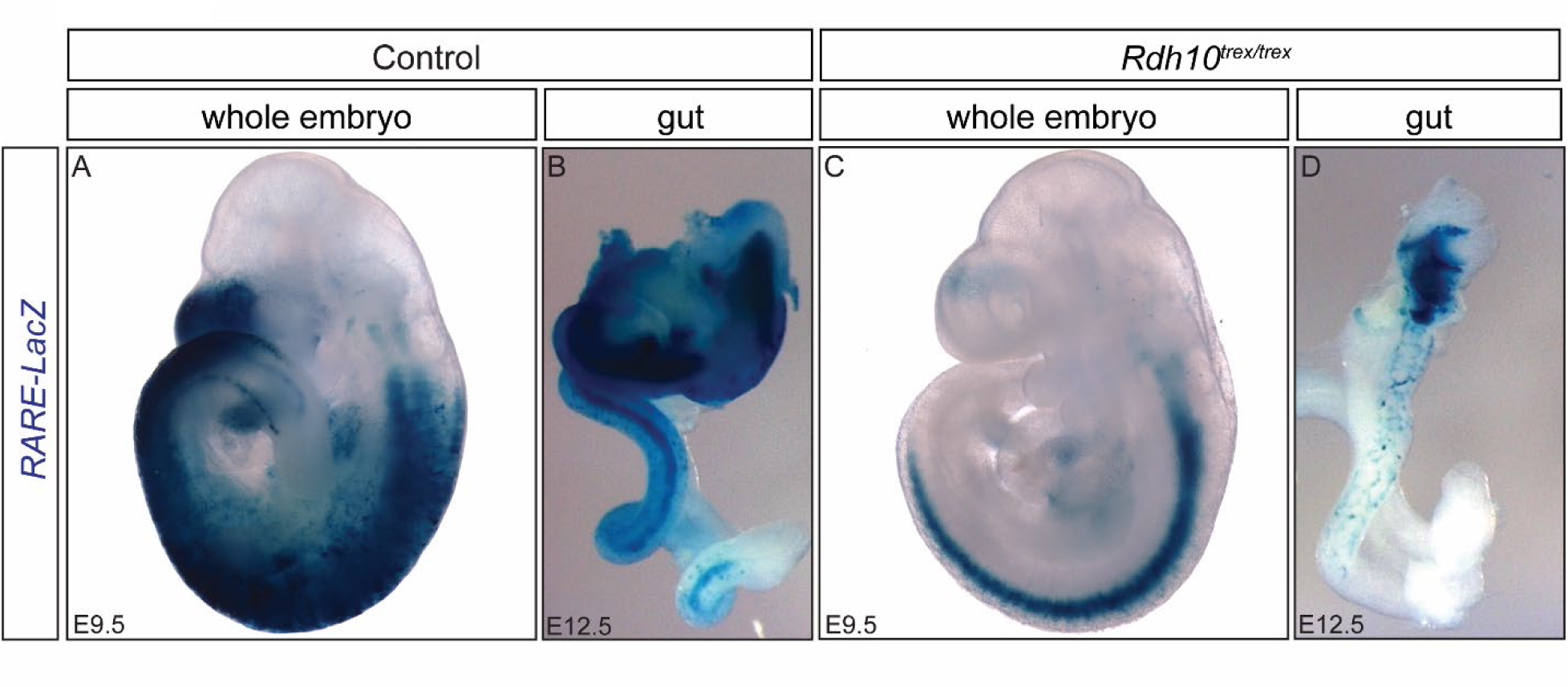
*Rdh10^trex/trex^* is retinoid signaling deficient. (A-D) The *RARE-lacZ* allele was bred into the background of *Rdh10^trex/trex^*embryos to visualize active RA signaling in blue via LacZ staining. At E9.5, (A) staining for RA signaling is present in the forebrain and hindbrain/spinal cord junction extending to the tail in control embryos. *Rdh10^trex/trex^*(C) shows greatly reduced signaling with staining restricted to the ventral neural tube. (B,D) Whole guts harvested from control and *Rdh10^trex/trex^* embryos at E12.5. (B) Control gastrointestinal tract shows broad LacZ staining for RA signaling in the stomach and intestine extending to the cecum. (D) *Rdh10^trex/trex^* gastrointestinal tract displays diminished LacZ staining in the stomach with nearly absent staining in the intestine.

In contrast, *Rdh10^trex/trex^* embryos exhibit a complete absence of RA signaling prior to E8.75 (10 somite stage)(19, 33, 34). E9.5 *Rdh10^trex/trex^* embryos display limited RA activity, which is confined to the ventral neural tube, and is absent from the adjacent ventral mesenchyme (Figure 3C). Of note, the developing cranium is nearly absent of RA signal at E9.5 in *Rdh10^trex/trex^; RARE-LacZ* embryos yet cranial neural crest cells have successfully migrated to distal branchial arches in the *Rdh10^trex/trex^* embryos (18, 32). Retinoic acid signal can be contrasted to the *Rdh10* spatiotemporal expression between E8.5-10.5 shown by LacZ staining of the *Rdh10^βgeo^*mouse line (Figure 1E-L). *Rdh10* expression is a smaller subset of active signal in the neural tube-somite-mesenchyme axis as compared to *RARE-lacZ*.

Diminished RA signaling in *Rdh10^trex/trex^* embryos is particularly evident in the gastrointestinal tract. In guts isolated from E12.5 control embryos, *RARE-LacZ* activity detected throughout the stomach and intestine extending distally to the cecum (Figure 3B). In contrast, in E12.5 *Rdh10^trex/trex^* guts, the domain of RA signaling is considerably reduced in the hypoplastic stomach, and is diminished in the small intestine (Figure 3D). Collectively, these data suggest that *Rdh10* loss-of-function results in reduced RA signaling in vagal level mesoderm and foregut mesenchyme, which spatiotemporally correlates with the perturbation of NCC migration and entry into the foregut.

### *Rdh10^trex/trex^* intestinal aganglionosis is not caused by delayed NCC migration

*Rdh10^trex/trex^* embryos typically die between E12.5-14.5, due to cardiovascular failure and hemorrhage. To bypass this lethality and evaluate NCC migration defects as a cause of intestinal aganglionosis, we isolated GI tracts from E11.5 control and *Rdh10^trex/trex^* embryos and cultured themin retinoid-deficient media for 2, 4 or 10 days (Supplemental Figure 3A-H). E11.5 control embryos have mature neurons in the primitive GI tract extending from the foregut to the cecum between the midgut and hindgut as evidenced by TUJ1 staining. In contrast, the GI tract in *Rdh10^trex/trex^* embryos exhibits neuronal agenesis (Figure 1 B, D; Supplemental Figure 3 A, E). In controls, after 2 days of culture, NCC have colonized the hindgut and begun to form a dense neural network. After 4 days and continuing to 10 days of culture, the intestines in control embryos exhibit a thick, reticulated neural network extending into the hindgut (Supplemental Figure 3B-D). In contrast, the intestines of *Rdh10^trex/trex^*embryos exhibit little to no neuronal staining (Supplemental Figure 3F,G,H) irrespective of the length of time they were cultured. These *in vitro* culture experiments suggest that the gut phenotype observed in E11.5 *Rdh10^trex/trex^* embryos is not due to delayed vagal NCC migration or defects in their survival, but rather a failure of NCC to enter the foregut in association with perturbed *Rdh10*-mediated RA signaling.

To further substantiate a requirement for RA signaling in NCC entry and colonization of the GI tract, we examined a second retinoid signaling deficient mouse line, *Aldh1a2^gri/gri^*, which we also recovered in our ENU mutagenesis screen(18). *Aldh1a2* functions in the second step of vitamin A metabolism, oxidizing retinal to retinoic acid and we previously showed *Aldh1a2^gri/gri^* embryos exhibit a near total absence of *RARE-LacZ* activity, consistent with a loss of RA signaling(19). *Aldh1a2^gri/gri^* embryos survive until approximately E10.0 and *Sox10* labeling of NCC revealed abnormal patterns of cranial and trunk NCC, including defective ventral migration of vagal NCC. The NCC in *Aldh1a2^gri/gri^* mutant embryos also fail to enter and colonize the GI tract in contrast to littermate controls (Supplemental Figure 3I-J’). Taken together with the phenotype of *Rdh10^trex/trex^* embryos, our data reveals that RA signaling is essential for vagal NCC invasion and colonization of the foregut.

### *Rdh10*-mediated retinoid signaling is required prior to E11.5 for NCC migration, entry, and colonization of the GI tract during ENS development

The association between the spatiotemporal activity of *Rdh10* and RA signaling during the migration and entry of vagal NCC into the foregut raised the possibility that the intestinal aganglionosis phenotype in *Rdh10^trex/trex^*embryos could be rescued via retinoid supplementation. Consistent with this idea, previous work has suggested that RA signaling promotes NCC migration throughout the GI tract(18). Therefore, we tested whether retinoid supplementation could rescue the migration of vagal NCC and their entry into the foregut and subsequently facilitate enteric neuron differentiation and ENS formation in *Rdh10^trex/trex^* embryos. Intestines isolated from E11.5 control and *Rdh10^trex/trex^* embryos were cultured with either all-trans-retinoic acid or retinal for 4 days each (Supplemental Figure 4A-F). Compared to untreated controls, wild-type GI tracts cultured with 25nM of all-trans-RA, a sub-physiological, non-teratogenic dose(22), continued to display thick, reticulated neural networks, as evidenced by TUJ1 immunostaining (Supplemental Figure 4A,B). In contrast, cultured *Rdh10^trex/trex^* intestines remain largely devoid of TUJ1 immunostaining indicative of persistent intestinal aganglionosis (Supplemental Figure 4D,E). All-trans-RA has the potential to cause teratogenic defects(18), therefore, to ensure a more relevant physiological effect, we supplemented cultured control and mutant GI tracts with 0.3nM(18) retinal since retinal will only be converted to RA in an appropriate temporal and tissue specific manner by *Aldh1a2* (*Raldh2*). Control GI tracts that were supplemented with retinal exhibited neural networks throughout the gut but were grossly less complex than in control untreated cultured intestines (Supplemental Figure 4A,C). In contrast, *Rdh10^trex/trex^* GI tracts remained aganglionic (Supplemental Figure 4F). Thus, culturing E11.5 *Rdh10^trex/trex^* gastrointestinal tracts for 4 days with either retinal or RA provides no improvement in colonization or neuronal differentiation. The results of our *in vitro* GI tract organ culture experiments confirm the absence of NCC from the foregut in *Rdh10^trex/trex^* embryos, and suggest retinoid signaling is required prior to E11.5 for proper NCC migration, entry and colonization of the GI tract, during ENS development.

### *Rdh10^trex^* intestinal aganglionosis can be rescued by *in utero* dietary retinal supplementation

To refine the temporal window during which retinoid signaling is required for NCC migration and ENS formation, we mated *Rdh10^trex/+^* mice together, and supplemented pregnant dams with retinal (12.5mg all-trans-retinal/kg maternal body weight) via oral gavage once per day from E7.5-E11.5 (Figure 4 A,B,D,E). Wild-type and mutant embryo GI tracts were subsequently harvested at E11.5 for analysis. Once daily retinal gavage of control embryos from E7.5-E11.5 revealed typical enteric neuron development with an extensive network throughout the stomach and intestine extending toward the cecum which was comparable to untreated controls (Figure 4 A,B). Unlike the aganglionic phenotype in untreated *Rdh10^trex/trex^*embryos (Figure 4D) retinal treated *Rdh10^trex/trex^* embryos developed mature neurons in the foregut and intestine, extending towards the cecum (Figure 4E). These data suggest retinoid signaling is required prior to E11.5 for NCC entry, colonization and differentiation in the GI tract.

**Figure 4.**
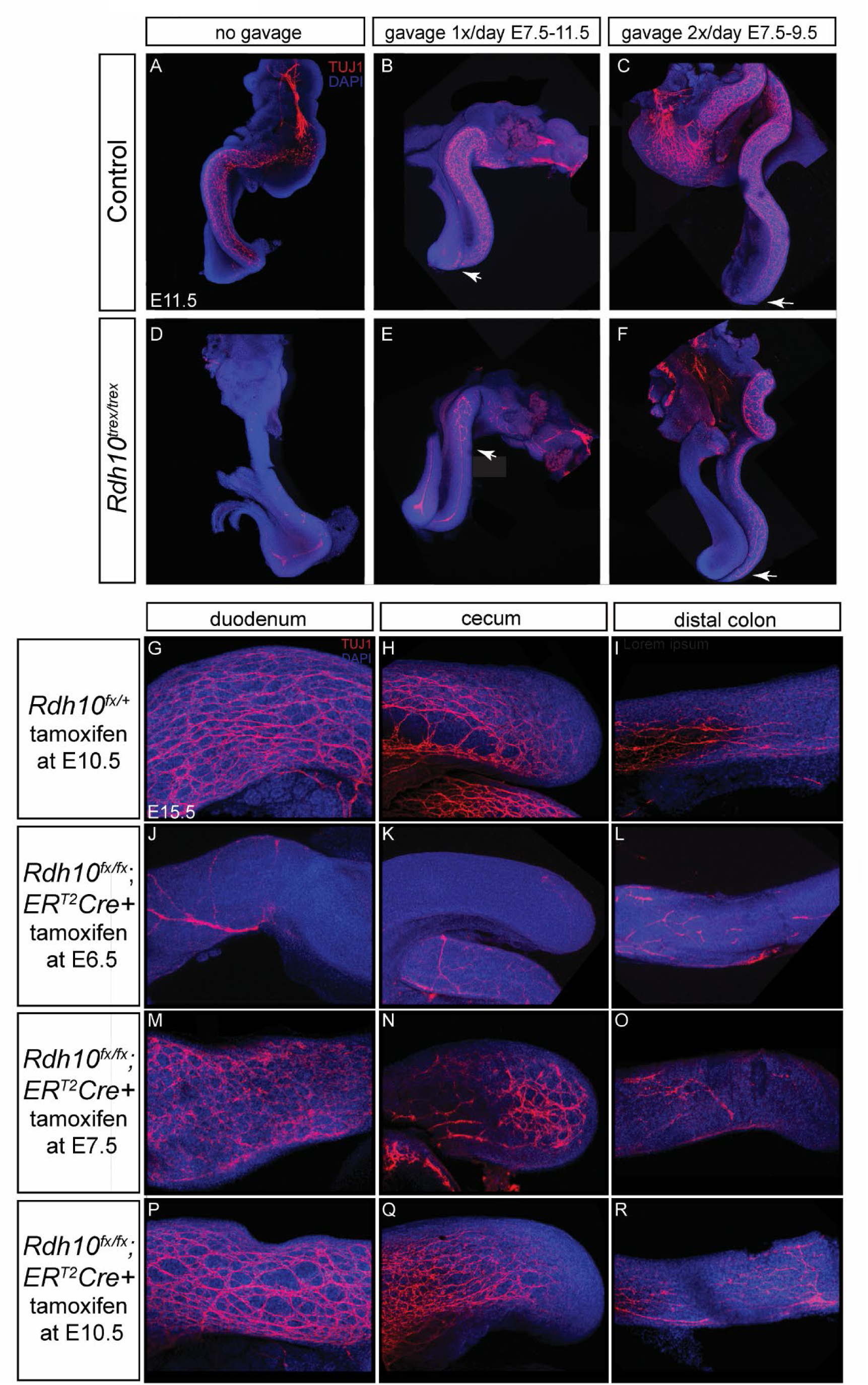
*Rdh10*-mediated RA signaling is required for early gut colonization. Retinal *in utero* gavage of pregnant *Rdh10^trex/+^* dams was used to rescue intestinal aganglionosis. (A-F) TUJ1 (red) and DAPI (blue) immunostaining in whole guts harvested at E11.5. (A,D) Non-gavaged control and mutant embryonic guts. Neurons are present in the control gut (A) in the stomach and midgut leading to the cecum. The mutant (D) lacks discernable neuronal staining entirely in the stomach, midgut and hindgut regions. (B,E) Pregnant dams were gavaged with retinal once daily between E7.5-11.5 with white arrows indicating the caudal most point of enteric neural development. The control whole gut (B) shows normal development with enteric neurons in the stomach and midgut leading to the cecum. *Rdh10^trex/trex^* gut (E) has neurons present in the stomach and the first third of the midgut. (C,F) Pregnant dams were gavaged with retinal twice daily between E7.5-9.5, which represents the minimal retinal treatment necessary to rescue the *Rdh10^trex/trex^* gut wavefront (C) to a comparable wild-type location (F) indicated by white arrows. (G-R) Temporal *Rdh10* deletion in *Rdh10^fx/fx^*; ERT2Cre by tamoxifen and progesterone gavage in utero at E6.5 (J-L), E7.5 (M-O) and E10.5 (P-R). Whole guts were harvested at E15.5 with TUJ1 (red) and DAPI (blue) immunostaining. Control littermates at each of these time points are represented by the *Rdh10^fx/+^* embryos given tamoxifen and progesterone gavage at E10.5 (G-I). Guts from embryos treated after E8.5 (P-R) appear indistinguishable from controls with full colonization and neural reticulation through the full gut. Tamoxifen treatment at E7.5 produces neural development in the duodenum (M) and cecum (N) with aganglionosis in the distal most portion of the colon (O), whereas treatment one day earlier at E6.5 results in aganglionosis in the duodenum (J), cecum (K) and colon (L).

To increase retinal availability in *Rdh10^trex/trex^* embryos, we performed twice daily supplementation with retinal on consecutive days beginning at E7.5. Embryos were harvested at E11.5. Compared with once daily retinal supplementation, twice daily supplementation between E7.5-E9.5, resulted in a phenotypic rescue of ENS formation in *Rdh10^trex/trex^*embryos by E11.5. Neuronal differentiation in *Rdh10^trex/trex^* GI tracts was comparable to treated and untreated control GI tracts, with mature neurons in the stomach and intestines, extending to the cecum (Figure 4 A,C,F). Collectively these results demonstrate *Rdh10* mediated RA signaling is required from E7.5 to E9.5 for proper ENS formation. This period immediately precedes vagal NCC entry into the foregut and colonization of the GI tract. These maternal gavage data imply that *Rdh10* is not required for the maintenance of NCC migration or colonization once NCC are already in the GI tract. Rather, *Rdh10*-mediated RA signaling is essential for vagal NCC migration and entry into the foregut. Consistent with these observations, we previously showed that retinal supplementation can rescue *Rdh10^trex/trex^* embryos through to birth with postnatal survival (22).

### Temporal deletion of *Rdh10* in *Rdh10^flox/flox^; ER^T2^Cre* mice confirms early requirement of retinoids during ENS development

To corroborate the spatiotemporal requirement for *Rdh10*-mediated RA signaling in ENS formation, we generated *Rdh10^flox/flox^* mice, which were crossed to a Tamoxifen inducible ubiquitous Cre line, *ER^T2^Cre*(*18*), which allows for the selective temporal excision of *Rdh10*, simultaneously overcoming the limitations of *Rdh10^trex/trex^* embryo lethality and the need for retinoid supplementation. Tamoxifen (5mg) and progesterone (1mg) were co-administered to pregnant dams via oral gavage at a single time point between E6.5 to E11.5.

*Rdh10^flox/flox^;ER^T2^Cre* embryos were subsequently harvested at E15.5 and assessed for NCC colonization of the gut and ENS differentiation (Figure 4G-R).

Temporal excision of *Rdh10* by E7.5, following Tamoxifen administration to *Rdh10^flox/flox^*; *ER^T2^Cre* embryos at E6.5, results in intestinal aganglionosis throughout the duodenum, cecum and distal colon (Figure 4J-L). In contrast, administration of Tamoxifen to *Rdh10^flox/flox^*; *ER^T2^Cre* embryos at E7.5 with subsequent deletion by E8.5 results in enteric neuron differentiation in the duodenum and cecum, but distal colonic aganglionosis, reminiscent of a short-segment Hirschsprung disease phenotype (Figure 4M-O). Temporal removal of *Rdh10* at later time points (E9.5 – E11.5) does not induce a HSCR-like phenotype (Figure 4P-R and data not shown). Normal gut colonization and ENS formation was observed in all treated *Rdh10^flox/+^* littermates irrespective of the age of treatment (E10.5 gavage shown Figure 4G-I). These conditional deletion studies further define the precise temporal requirement for *Rdh10*-mediated RA production and augments the data obtained from retinal-supplemented *in vivo* cultures of *Rdh10^trex/trex^* GI tracts. *Rdh10*-mediated RA signaling is therefore required between E7.5-9.5 during which vagal NCC are born, migrate ventrally and invade the foregut. In contrast, *Rdh10* appears not necessary for the maintenance of NCC migration and colonization of the GI tract after NCC have entered the foregut.

### Non-cell autonomous paracrine effect of *Rdh10*-mediated RA signaling on NCC

The early temporal requirement for *Rdh10-*mediated RA signaling to form a complete enteric nervous system raised the question of whether *Rdh10* is cell autonomously and intrinsically required in NCCs, or whether it functions non-cell autonomously in a paracrine fashion within the NCC microenvironment. To test for an intrinsic cell autonomous role in NCC, we conditionally deleted *Rdh10* in the dorsal neuroepithelium, the territory from which NCCs are derived using *Wnt1Cre*. E10.5 *Rdh10^flox/flox^;Wnt1Cre* and *Rdh10^flox/flox^* whole embryos exhibit normal peripheral nervous system development with clear neuronal labeling in the anterior GI tract (Supplemental Figure 5A,B). Whole GI tracts isolated from E18.5 *Rdh10^flox/flox^* as well as *Rdh10^flox/flox^;Wnt1Cre* embryos displayed a similar gross neuronal network and density along the entire length of the GI tract (Supplemental Figure 5C-H). Furthermore, *Rdh10^flox/flox^;Wnt1Cre* mice are viable postnatally and do not exhibit any obvious motility or gastrointestinal dysfunction as measured by carmine red whole gut transit assay. Because vagal nerve development and ENS formation were unaffected in NCC-specific conditional knockouts of *Rdh10*, *Rdh10* is therefore not intrinsically or cell autonomously required in vagal NCC for their migration and entry into the foregut, nor for their migration throughout the GI tract, or differentiation, during formation of the ENS.

### Targeted qRT-PCR and *in situ* hybridization reveal disruptions in *Ret* signaling axis in *Rdh10^trex/trex^* embryos

Mutations in the *RET-GDNF-GFRA1* signaling axis genes are principally associated with the etiology of HSCR. Interestingly, loss-of-function mouse models of each gene exhibit intestinal aganglionosis(35, 36), similar to the phenotype described above in *Rdh10^trex/trex^*embryos. Ret and Gfrα1 are co-receptors for Gdnf, and each play important roles in promoting NCC migration(37, 38) and ENS formation. *Ret* is expressed by migrating vagal neural crest cells and its expression is maintained throughout colonization of the gut(39). Similarly, *Gfrα1* is also expressed by NCC. *Gdn*f however, is expressed primarily in the mesenchyme where it functions in a paracrine manner as a chemoattractant for NCC to invade the foregut. We used qRT-PCR and *in situ* hybridization to assess *Rdh10* mediated regulation of this pathway by comparing the levels and spatial domains of gene expression. qRT-PCR revealed that the levels of *Ret, Gdnf* and *Gfrα1* mRNA in E9.5 *Rdh10^trex/trex^* embryos were significantly reduced to 54%, 70% and 49% (p<0.05), respectively of wild-type levels (Figure 5A). We verified the decrease in *Ret* and *Gdnf* expression by *in situ* hybridization (Figure 5 B-I). In E9.5 control embryos, *Ret* expression was evident within the foregut extending caudally to the midgut (Figure 5 B,B’). In contrast, there was a complete absence of *Ret* labeled cells in the developing gut of *Rdh10^trex/trex^* mutant embryos (Figure 5 C,C’). To evaluate if a disruption in ligand signaling underlies the mutant phenotype, we evaluated the activity of the Ret ligand, Gdnf, which is normally expressed within the foregut mesenchyme through which vagal neural crest cells migrate in E9.25-9.5 wildtype embryos (Figure 5 D, D’, F, F’). We found that Gdnf expression was decreased in *Rdh10^trex/trex^* mutant embryos (Figure 5 E,E’G,G’). Sections of E9.5 control embryos highlight the expression of *Gdnf* within the foregut mesenchyme and epithelium (Figure 5 H,H’). In contrast, *Rdh10*^trex/trex^ embryos exhibit reduced levels of *Gdnf* in the foregut mesenchyme and epithelium (Figure 5 I, I’), consistent with the qRT-PCR results (Figure 5 A).

**Figure 5.**
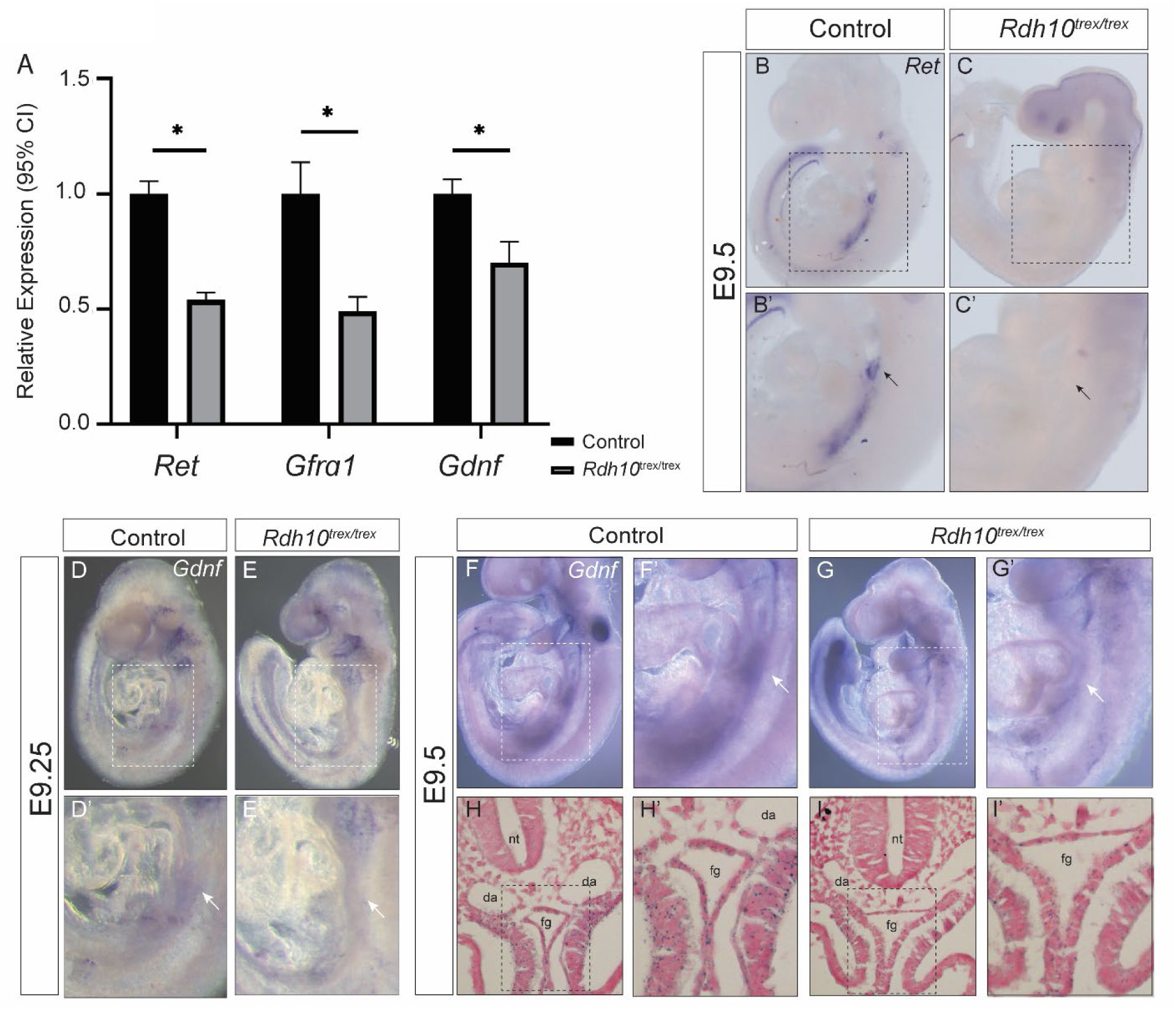
*RET-GDNF-GFRA1* signaling axis signaling is disrupted in *Rdh10^trex/trex^*. A. qRT-PCR results show that *Rdh10^trex/trex^*embryos have a significantly reduced level of *Gdnf, Ret* and *Gfrα1* compared to control embryos. Relative quantities of each gene are graphed with 95% confidence intervals. Each of these three genes expression level differences showed significant p-values (p<0.05), as indicated by asterisk. *Gdnf* p=0.0168, *Ret* p=1.93E-6, *Gfrα1* p= 0.0011. B-E. *In situ* hybridization was used to assess *Ret* and *Gdnf* expression domains. B, B’. E9.5 Control embryos express *Ret* in the foregut and mid-gut shown by the black arrowhead. C, C’. *Rdh10*^trex/trex^ embryos completely lack *Ret*-positive staining within the gut. D, D’, F, F’. Control E9.25 and E9.5 embryos express *Gdnf* within the foregut entrance indicated by the white arrows. E, E’, G, G’. *Rdh10*^trex/trex^ embryos, however, have a significant downregulation of *Gdnf* within the foregut shown by the white arrow. H, H’, I, I’. Sections of E9.5 control embryos of foregut mesenchyme and epithelium with more *Gdnf* signal present in control than *Rdh10*^trex/trex^, consistent with the qRT-PCR results (A).

To further evaluate if there was an association between RA signaling and *Gdnf* activity, we examined the expression of *Gdnf* in E9.5 *Aldh1a2^gri/gri^* embryos, which we previously showed were deficient in RA signaling(19) similar to *Rdh10^trex/trex^*embryos. E9.5 control embryos display *Gdnf* expression within the foregut (Supplemental Figure 6 A,A’, C, C’). In contrast, *Aldh1a2^gri/gri^* embryos exhibit diminished *Gdnf* activity within the foregut region (Supplemental Figure 6 B,B’, D, D’). The downregulation of *Gdnf* expression in *Aldh1a2^gri/gri^* embryos is consistent with our results in *Rdh10^trex/trex^* embryos and substantiates a role for RA signaling in regulating *Gdnf* expression during NCC entry into the foregut. Since *Rdh10* is not intrinsically required in vagal NCC, our qRT-PCR and expression analyses in *Rdh10^trex/trex^* and expression data in *Aldh1a2^gri/gri^*embryos indicate that *Rdh10* operates in a paracrine fashion to promote vagal NCC migration and entry into the foregut in part by functioning upstream of *Ret-Gdnf-Gfrα1* signaling.

### RNA sequencing reveals alterations in ENS development genes in *Rdh10*^trex/trex^ embryos

Our data indicates that RA signaling plays a critical role in regulating the NCC microenvironment. To gain a deeper understanding of the impact of RA signaling, we performed comparative transcriptomics analyses via bulk RNA-sequencing of the vagal region in E9.5 control and *Rdh10^trex/trex^* littermate embryos. RNA sequencing identified 20,883 genes that were then analyzed by EdgeR, which revealed 4,366 genes that were differentially regulated between *Rdh10*^trex/trex^ and control embryos with a p-value < 0.05. Of these genes, 1,095 were changed by at least 1.74-fold with a p-value < 0.05, including 533 upregulated and 562 downregulated genes (Figure 6). With this gene set, we used targeted analysis to identify genes that have a known role in ENS development and HSCR etiology, as well as an unbiased approach using enrichment analysis to uncover critical genes and pathways that are regulated by RA signaling during ENS formation.

**Figure 6.**
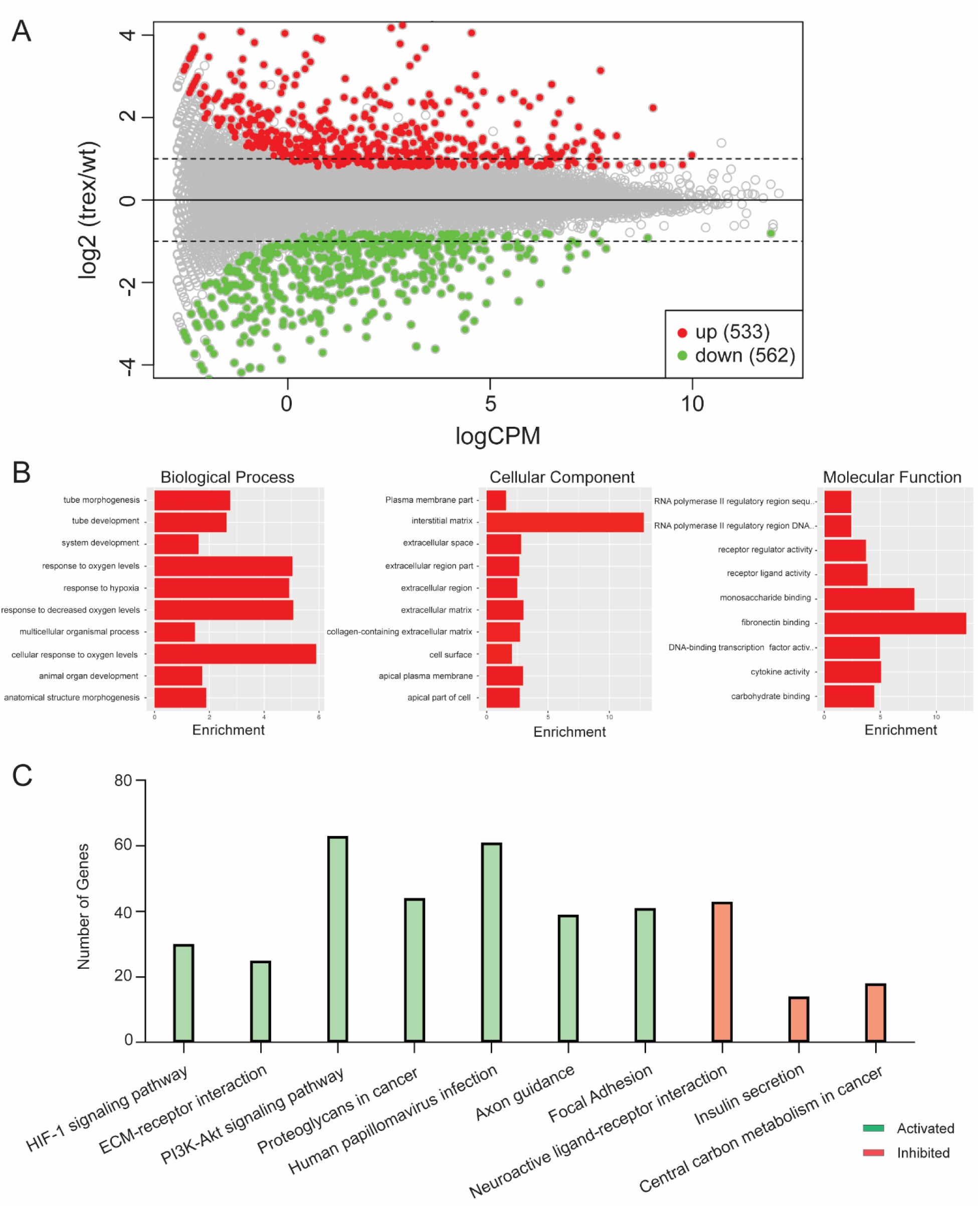
RNA sequencing revealed numerous ECM pathway targets. A. MA plot for RNA sequencing. Genes with p-values less than 0.01 and a fold change greater than 1.74-fold are highlighted in red (upregulated in *Rdh10*^trex/trex^) or green (downregulated in *Rdh10*^trex/trex^). B. The top up and down regulated genes that changed by at least 1.74-fold with a p-value < 0.05 were examined for enrichment of gene ontology (GO) terms. This identified enrichment in several biological processes related to cellular responses to oxygen levels, decreased oxygen levels and hypoxia. Cellular processes related to interstitial and extracellular matrices, and Molecular functions related to fibronectin binding. C. Signaling Pathway Impact Analysis (SPIA) algorithm was used to assess Kyoto Encyclopedia of Genes and Genomes (KEGG) pathway perturbations of all genes with adjusted p-value < 0.05.

First, we used a targeted analytical approach to identify genes that had ≥ 2-fold expression change with a p-value < 0.05, and that when mutated were already known to produce an overt intestinal phenotype in mouse models. *Gfra1, Phox2b and Ecel1* were downregulated, whereas *Col6a4* was upregulated in *Rdh10^trex/trex^* embryos compared to controls (Supplemental Table 1). *Gfrα1* is a receptor for *Gdnf* and therefore a major regulator of the *Ret-Gdnf-Gfrα1* signaling axis as is *Phox2b,* which can specifically regulate *Ret* expression(40)*. Phox2b* downregulation was validated using immunohistochemistry (Figure 7 A-G). *Gfrα1* and *Phox2b* loss-of-function mouse mutants exhibit intestinal aganglionosis(41–43), whereas *Ecel1* mutants present with distal colonic aganglionosis(*44*), consistent with each of these genes functioning in vagal NCC migration and ENS formation(43, 44). Furthermore, mutations in *GFRA1, PHOX2B* and *ECEL1* have all been identified in patients with HSCR(45–47). The same is also true for *PAX3 and EDNRB*, and loss-of-function mouse models exhibit total (*Pax3*) or distal (*Ednrb*) intestinal aganglionosis. Both *Pax3* and *Ednrb* were also downregulated in *Rdh10^trex/trex^*embryos compared to controls. In addition, we noted downregulation of specific genes known to regulate *Ret* expression such as *Rarβ, Hoxb5 and Nkx2.1*(*40, 48*) (p-value < 0.05) (Supplemental Table 1). *RARβ* is a receptor for RA, is expressed in NCC, and binds to enhancer regions of *Ret* to promote its activity(49). *Hoxb5* is expressed in vagal NCC and functions in ENS formation as *Hoxb5* mouse mutants exhibit hypoganglionosis(50). *Hoxb5* also forms a complex with *Nkx2.1* to synergistically regulate *Ret*, as well as coordinate with *Phox2b* and *Pax3* to mediate expression at the *Ret* promoter(40, 50). In addition to *Hoxb5*, we identified several other *Hox* genes that were also differentially expressed by over 2-fold with a p-value < 0.05 (Supplemental Table 1). This is consistent with *Hox* gene activation by RA during anterior-posterior patterning of the gut and their association with ENS formation and HSCR pathogenesis (*51–53*).

**Figure 7.**
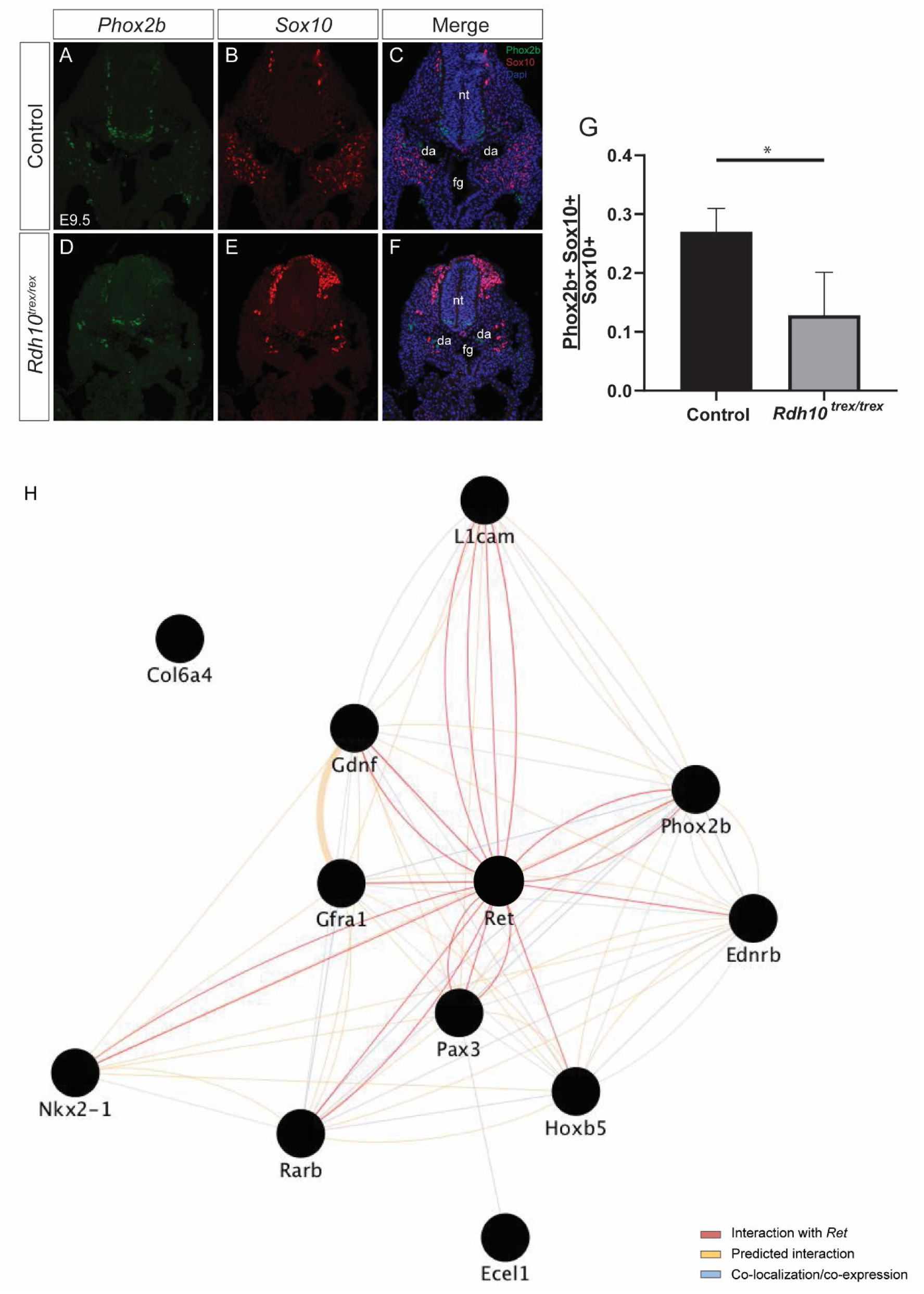
ENS protein expression of *Phox2b* and ENS network regulated by *Rdh10*-mediated RA signaling. A-F. E9.5 embryos co-stained for *Phox2b* and *Sox10. Phox2b* stains enteric precursor cells, and *Sox10* marks differentiated neurons. A-C Control embryos stained for Phox2b, Sox10 and Merge. D-F *Rdh10*^trex/trex^. G. The ratio of NCC to neurons is significantly decreased in *Rdh10*^trex/trex.^ H. RNA sequencing identified a series of genes that are dysregulated in *Rdh10^trex/trex^* that have critical roles in ENS formation and HSCR pathogenesis. The interaction between these genes were mapped by Cytoscape to show gene interactions that include, interactions with *Ret* (red lines), predicated interactions between genes (yellow lines) and genes in literature that are either co-expressed or co-localize (blue lines).

In contrast to downregulation of the genes noted above, others such as, *Col6a4*, were upregulated in *Rdh10*^trex/trex^ embryos (Supplemental Table 1). Collagen VI immunostaining appeared increased in *Rdh10*^trex/trex^, but was not statistically significant when quantified as fluorescent intensity (Figure 8 E-H, L), However, *Col6α4* is expressed in the extracellular matrix of the developing gut and known to negatively regulate NCC migration(54). In fact, increased *Col6a4* production can cause distal colonic aganglionosis in mouse models(54) and patients with HSCR have been diagnosed with abnormally high levels of Collagen VI surrounding the myenteric plexus(54). The misregulation of genes involved in ENS development, *Ret* signaling and the etiology of HSCR demonstrate that *Rdh10* mediated RA signaling regulates a diverse genetic network that is instrumental in promoting vagal NCC development during ENS formation and in the pathogenesis of HSCR (Figure 7H).

**Figure 8.**
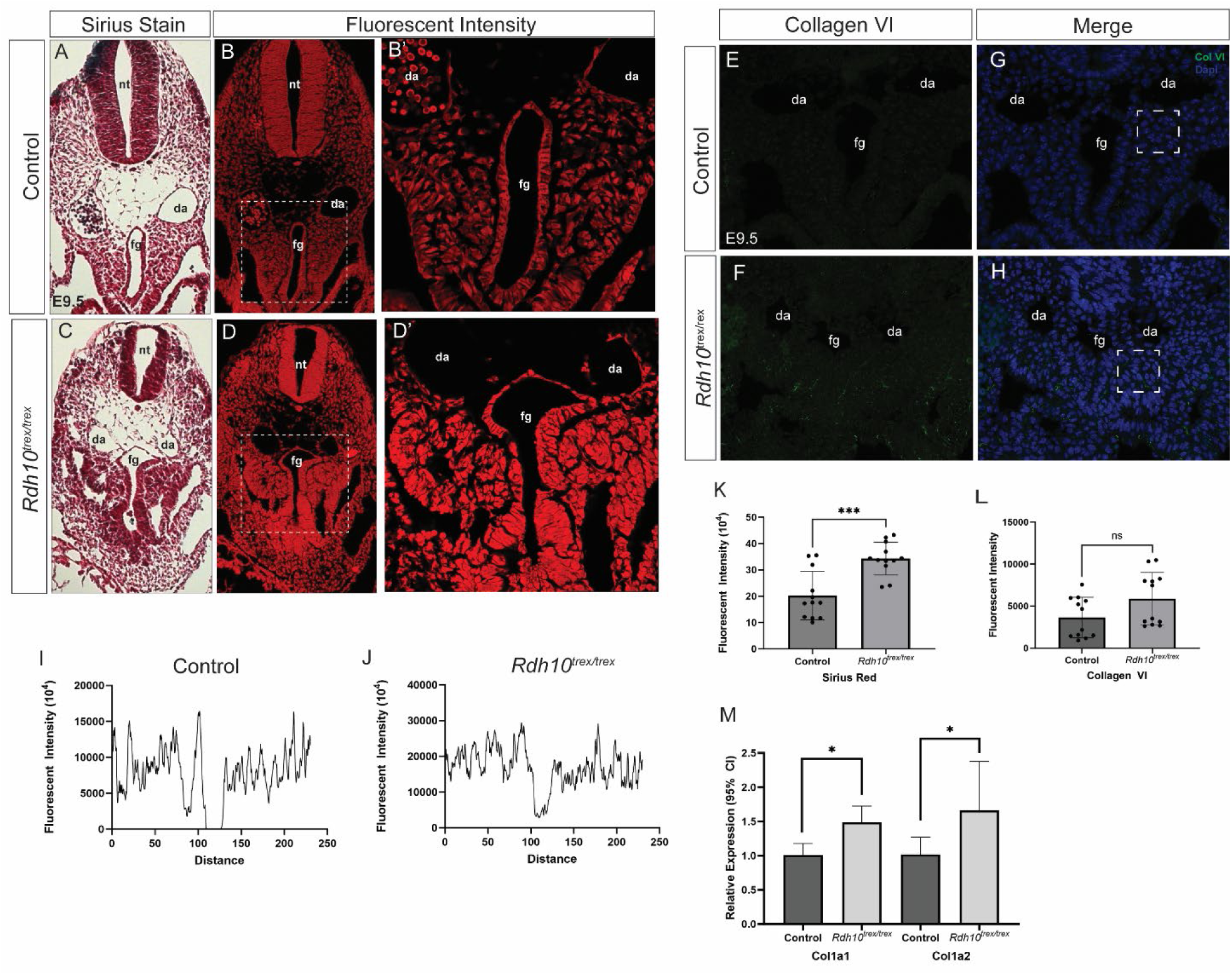
Increased collagen in E9.5 *Rdh10*^trex/trex^ compared to control embryos. Sirius red staining and fluorescent intensity show in control embryos (A, B, B’). *Rdh10*^trex/trex^ embryos (C, D, D’). Inset B’ and D’ is quantified in I, J and compared in K. Significant (P<0.05) increase in collagen as measured by Sirius red fluorescent intensity in *Rdh10*^trex/trex^. Collagen VI staining for E 9.5 Control (E, G) versus *Rdh10*^trex/trex^ (F, H). Collagen VI immunostaining appeared increased in *Rdh10*^trex/trex^, but was not statistically significant when quantified as flurorescent intesity (Figure 8 L). M. qRT-PCR of Collagen 1a1 and 1a2 significantly increased in *Rdh10*^trex/trex^. P<0.05.

### RNA sequencing reveals significant alterations in extracellular matrix proteins and focal adhesion proteins in *Rdh10^trex/trex^*embryos

We utilized RNA sequencing enrichment analyses to identify gene and pathway changes in an unbiased manner. The relationship between gene expression ratio (trex/wt) and signal strength is displayed in the MA plot. Genes with p-values less than 0.01 and a fold change greater than 1.74-fold are highlighted in color (Figure 6A). The top up and down regulated genes were examined for enrichment of gene ontology (GO) terms. This identified enrichment in several biological processes related to cellular responses to oxygen levels, decreased oxygen levels and hypoxia (Figure 6B). Elevated hypoxia is associated with defects in vasculogenesis(55). This is notable as RA regulates endothelial cells during vascular modeling and patterning(56). Retinoid deficient models of *Raldh2* have defects in vasculogenesis in the yolk sac(56) and cranial vascular plexi(18). Similarly, *Rdh10^trex/trex^* embryos display defective vasculogenesis as evidenced by a diminished dorsal aorta(18) and hemorrhaging that likely contributes to embryo lethality(19). In addition to these biological processes, we identified enrichment in the cellular components of interstitial and extracellular matrix, and extracellular space (Figure 6 B). Interestingly, the interstitial matrix is a specific type of extracellular matrix found in interstitial connective tissue, characterized by the presence of fibronectins, and types I, III, V, VI, VII and XII collagens, many of which were upregulated in *Rdh10^trex/trex^* embryos (Figure 6, Supplemental Table 2). Fibronectin is localized to the extracellular matrix and promotes NCC migration(57–60), in addition to binding other components of the interstitial and extracellular matrix, including collagens(60–62). However, Collagen VI can impede fibronectin’s positive affect on NCC migration(54, 63), illustrating the importance of matrix composition and its effects.

In addition to GO analysis, we performed Signaling Pathway Impact Analysis (SPIA)(64) to assess Kyoto Encyclopedia of Genes and Genomes (KEGG) pathway perturbations on all genes with an adjusted p-value of 0.05 or less. This returned 1992 genes, of which 1851 mapped to Entrez IDs. We noted an enrichment in ECM-receptor interaction and focal adhesion pathways in *Rdh10*^trex/trex^ embryos (Figure 6B), which are also known to affect NCC migration. To validate the findings using SPIA, we employed additional pathway enrichment programs, such as ToppGene and DAVID, which consistently revealed enrichment in matrix and adhesion. Some of these pathways included ECM organization and interactions, focal adhesion, and collagen biosynthesis (Supplemental Table 2). Analysis by ToppGene also identified enrichment in Hif1α signaling, which was the top pathway identified by SPIA (Figure 6C; Supplemental Table 2). We noted GO term enrichment in several biological processes related to hypoxia, and *Hif1α* is a hypoxia-inducible factor that regulates signaling targets in low oxygen conditions. The pathways identified by SPIA, DAVID and ToppGene for ECM receptor interaction and focal adhesion, also align with our GO analysis as many of their gene products localize to the interstitial and extracellular matrix, and space. Additionally, receptor ligand activity identified by GO analysis corresponds in molecular function to many of the genes in the ECM receptor interaction pathway (Figure 6; Supplemental Table 2).

To identify gene changes related to ECM receptor interaction and focal adhesion, we utilized Ingenuity Pathway Analysis (IPA) which identified 128 differentially regulated genes (p-value ≤ 0.05) associated with ECM organization, binding, or adhesion, including several known to affect NCC migration (Supplemental Table 3). For example, the adhesion molecule *L1cam*, which was downregulated, promotes chain migration of NCC and is a modifier of ENS formation(65, 66). In contrast, *Laminin B1*, which was increased, is inhibitory to NCC migration in the gut(67). Furthermore, there was a sizeable upregulation in numerous distinct collagens which aligned with GO term enrichment in collagen-containing interstitial and extracellular matrix (Figure 6A) as well as DAVID pathway analysis enrichment in collagen biosynthesis and modifying enzymes (Supplemental Table 2).

We validated the increase in collagen using both qRT-PCR and Sirius red staining (Figure 8). qRT-PCR confirmed that both *Col1a1* and *Col1a2* were significantly upregulated in *Rdh10*^trex/trex^ with a p-value < 0.05 (Figure 8 A). The regulation of collagen by RA is not well understood, however RA is known to directly regulate the *Col1a1* gene(68), as well as *Col1a2* through direct binding of its promoter by *Rarβ*(69, 70). Picro-Sirius red dyes specifically label components of collagen(71), and we used Sirius red staining to evaluate global alterations in collagen composition in the extracellular matrix. In comparison to Sirius red stained sections of E9.5 wild-type embryos, we observed a significant (p<0.05) increase in collagen throughout the foregut of *Rdh10*^trex/trex^ embryos (Figure 8 A-D’). Greater than 50% increase was quantified by measuring the fluorescent intensity of Sirius red staining within a defined region of the anterior foregut mesenchyme which coincides with the entry of neural crest cells into the gastrointestinal tract (Figure 8 B’, D’, I-K). The upregulation of collagen was notable as individual components of collagen such as, *Col6a4*(*54*), as well as collectively high levels of collagen in the gut can reduce NCC migration resulting in abnormal intestinal development(72). Increased collagen composition in the extracellular matrix has previously been associated with increased stiffness and the inhibition of cell migration(54). Therefore, the increase in collagen deposition may increase matrix stiffness in the anterior foregut mesenchyme in *Rdh10*^trex/trex^ embryos, which in turn impedes NCC migration. In concert with decreased *Ret-GDNF-GFRa1* signaling, this could restrict the entry of vagal NCC into the foregut, resulting in their failure to colonize the gastrointestinal tract and form the ENS.

## Discussion

### *Rdh10* and RA signaling are required prior to vagal NCC invasion of the foregut during ENS formation

Retinoic acid (RA) has been implicated as an important morphogen in ENS development(73–76) and is temporally required for enteric neural crest cell chain migration into the foregut(77). Most mammalian models of RA deficiency however required dietary modification to reveal an intestinal phenotype. For example, mice with genetic deletion of RBP exhibit low retinol concentrations in their circulation(78) and give birth to viable offspring(79). Only when challenged with dietary vitamin A deprivation do *Rbp4*^-/-^ mutant mice elicit distal colonic aganglionosis(75).. Conversely, *Raldh2*^-/-^ mutant embryos are lethal around E8.5 prior to ENS formation(26), but if maternally supplemented with RA, they survive long enough to present with intestinal aganglionosis and other developmental anomalies(80). Although *in vitro* and *in vivo* studies revealed a link between *Rbp4* and *Raldh2* (*Aldh1a2*)(75, 76, 80–82), the manifestation of an intestinal phenotype only occurred in association with dietary modification, leaving unresolved the spatiotemporal requirement and function of RA during NCC invasion and colonization of the gut. A recent study defined cell-autonomous roles for the RA receptor α (*RARα*) using numerous Cre-drivers and found that when activating the dominant negative *RarαDN* in premigratory and migratory stages, mice develop intestinal aganglionosis, but conditional deletion at later stages result in much milder phenotype(83). In human HSCR bowel tissue, RARα and its coregulator CREB-binding protein (CBP) were expressed in ganglionated bowel segments but were reduced in aganglionic segments(84). We discovered that *Rdh10^trex/trex^* embryos, which are RA signaling deficient, survive until E13.5-14.5, and present with intestinal aganglionosis. This characteristic makes *Rdh10^trex/trex^* embryos a useful genetic model to study the effects of vitamin A metabolism on vagal NCC development and ENS formation.

Using an allelic series of *Rdh10* mice, we determined that Rdh10-mediated RA signaling is required in a paracrine fashion between E7.5-9.5 for vagal NCC invasion of the foregut. This early requirement was revealed by deleting *Rdh10* in a temporal manner throughout the embryo with *ER^T2^Cre.* Excision between E7.5-9.5 resulted in intestinal aganglionosis (Figure 4). In contrast, excision after E9.5 did not impact ENS formation, indicating that once NCC are within the foregut, *Rdh10* does not appear to be required for their continued migration and, or differentiation throughout the rest of the GI tract. Consistent with this idea, twice daily gavage with retinal between E7.5 to E9.5 was sufficient to rescue intestinal aganglionosis in *Rdh10^trex/trex^* embryos (Figure 4). Our studies therefore in combination with prior work in zebrafish, xenopus, avian, mice and humans suggests a conserved requirement for RA signaling in ENS development.

### *Rdh10* and RA signaling are required to facilitate vagal NCC migration into the foregut entrance

HSCR associated intestinal aganglionosis has previously been attributed to perturbed NCC migration or differentiation throughout the gut (85, 86). Therefore, to assess the mechanistic role of *Rdh10*-mediated RA signaling, we indelibly labeled NCC and their descendants in *Rdh10^trex/trex^* embryos and observed that they fail to invade the foregut (Figure 2). Interestingly, *Rdh10* is highly expressed in the mesenchyme through which the vagal NCC migrate (Figure 1) and RA signaling is diminished in this territory in *Rdh10^trex/trex^* embryos. We did not observe apoptosis specific to the vagal NCC population (Supplemental Figure 2), nor did we observe gastrointestinal defects when *Rdh10* was conditionally deleted in neural crest cells (Supplemental Figure 5). Thus, the inability of vagal NCC to migrate into the foregut in *Rdh10^trex/trex^*embryos points to a role for *Rdh10*-mediated RA signaling as a paracrine factor in the NCC microenvironment. In further support of this idea, we showed that *Aldh1a2^gri/gri^* embryos which are also deficient in RA signaling, exhibit a similar failure in NCC invasion of the foregut (Supplemental Figure 3). Therefore, Rdh10-mediated spatiotemporal RA signaling facilitates vagal NCC invasion of the foregut, which is a critical step in NCC migration and colonization of the GI tract during ENS formation.

### ENCC developmental and microenvironment genes are altered in *Rdh10^trex/trex^* embryos

Mutations in *RET*, *GDNF* and *GFRA1* are principally associated with the etiology of HSCR, of which variable intestinal aganglionosis is a defining feature (87),(88). In fact, approximately 63% of individuals with HSCR possess mutations in the *RET* signaling gene network which result in an overall decrease in RET signaling(88). *Ret* and its co-receptor *Gfrα1* are expressed on the surface of vagal NCC, whereas their ligand, *Gdnf*, is expressed in the mesenchymal microenvironment through which vagal NCC migrate. This creates a ligand-receptor gradient important for the chemotactic entry of vagal NCC into the foregut around E9.5(37), and then again during their continued migration and colonization of the GI tract. *Ret*, *Gdnf* and *Gfra1* null mouse mutants therefore each exhibit intestinal aganglionosis(39, 42, 43, 89–91), and we discovered that expression of the *Ret* signaling network is downregulated in *Rdh10*^trex/trex^ embryos consistent with the pathogenesis of intestinal aganglionosis in *Rdh10*^trex/trex^ embryos. More specifically, *Ret* expression was absent in E9.5 *Rdh10*^trex/trex^ embryos and *Gdnf* expression was considerably diminished (Figure 5). Furthermore, *Gdnf* expression was also affected in RA signaling deficient *Aldh1a2^gri/gri^* embryos (Supplemental Figure 6). Taken together, this data is consistent with RA signaling being a positive regulator of the *Ret-Gfra1-Gdnf* genetic network and critical for NCC entry and colonization of the GI tract.

Interestingly, the *Ret* gene contains a putative retinoic acid response element (RARE) that binds retinoic acid receptors in the promoter/enhancer region upstream of the *Ret* transcriptional start site(92). We found using RNA sequencing data that *Rarβ* was downregulated over 2-fold in *Rdh10*^trex/trex^ embryos (Supplemental Table 1). Furthermore, a cis-regulatory element variant exists in humans within the enhancer region of *RET* that disrupts the binding of RARβ, thereby reducing *RET* expression(49). The presence of this variant increases the risk of HSCR in the human population by 1.7-fold and collectively suggests that RA directly regulates *RET* during normal ENS formation and in the pathogenesis of HSCR. Consistent with this idea, constitutive expression of a dominant-negative RAR, which blocks RA signaling, in NCC, diminishes Ret signaling in the ENS, resulting in aganglionosis (83). Conversely, exogenous RA can upregulate *Ret* in vagal NCC and promote their migration into the foregut (76).The upregulation of *Ret* by RA has been associated with increased tyrosine phosphorylation of Ret by Gdnf, implying RA can modulate Gdnf responsiveness(93).(83). Conversely, exogenous RA can upregulate *Ret* in vagal NCC and promote their migration into the foregut (76).The upregulation of *Ret* by RA has been associated with increased tyrosine phosphorylation of Ret by Gdnf, implying RA can modulate Gdnf responsiveness(93). Further evidence for an interaction between RA and other *Ret* pathway genes is evident from studies in which neurospheres treated with both RA and Gdnf, exhibited increased migration and increased neuronal connections(94). Moreover, RA also stimulates both *Gdnf* and *Gfrα1* expression *in vitro*(93, 95). Taken together, our data places *Rdh10*-mediated RA signaling upstream of the *Ret-Gfra1-Gdnf* signaling axis during vagal NCC entry into the foregut, which is essential for ENS formation, and when disrupted contributes to the pathogenesis of intestinal aganglionosis.

Although downregulation of *Ret-Gdnf-Gfrα1* expression could alone account for the intestinal aganglionosis phenotype in *Rdh10*^trex/trex^ embryos, our comparative RNA sequencing revealed misregulation of numerous other genes associated with NCC, ENS formation and HSCR etiology, including *Gfrα1, Phox2b*, *Pax3*, *Hoxb5*, *Ednrb, Ecel1,* and *Col6a4* (Figure 7, Supplemental Table 1). This aligns with a recent report that *in vitro* RA treatment is sufficient to induce expression of *Hoxb5*, *Phox2b* and *Pax3* (96). Interestingly, the role of these genes in HSCR pathogenesis has been linked to regulating *Ret*(40, 41, 48, 97, 98). Gfrα1 as a co-receptor for Gdnf, is a major component of the *Ret-Gdnf-Gfrα1* signaling axis. *Phox2b* null mutant embryos lack *Ret* in migrating ENCC(41) and *Phox2b* has been shown to independently regulate *Ret* expression *in vitro*(*40*). Both *Phox2b* and *Pax3* null mutants display total intestinal aganglionosis and mutations in *PHOX2B* have been reported in human cases of HSCR(41, 47, 97, 99–101). Pax3 interacts with Sox10 to bind an enhancer region of the *Ret* promoter and regulate its expression(97, 98). *Hoxb5* mutant mice present with an intestinal hypoganglionosis phenotype(50) and furthermore, Hoxb5 forms a complex with Nkx2.1, which synergistically regulates *Ret* activity, as well as, coordinates with Phox2b and Pax3 to mediate expression at the *Ret* promoter(40, 50).

The Endothelin pathway which also plays a central role in regulating vagal NCC migration and development in the gut(102) is disrupted in *Rdh10^trex/trex^* embryos, as evidenced by decreased expression of *Ednrb* and *Ecel1*. *Ednrb* and *Ecel1* loss-of-function mouse mutants exhibit distal colonic aganglionosis, and *Ednrb* mutants also display pigmentation defects, which together with HSCR phenocopies Waardenburg-syndrome in humans(103, 104). Variants of *Ednrb* increase the risk of HSCR(105, 106) and *Ednrb* interactions with *Sox10* enhance the penetrance and severity of HSCR phenotypes. Consistent with these findings, compound *Ednrb^sl/sl^*;*Sox10^Dom/+^*(**107**) double mutants exhibit total intestinal aganglionosis. Collectively these genes, identified by our comparative RNA sequencing, interact extensively in ENS formation and HSCR pathogenesis (Figure 7). Taken together with our data, this suggests that *Rdh10*-mediated RA signaling acts upstream of multiple signaling networks that coordinate vagal NCC migration and entry into the foregut during ENS formation and that are associated with the etiology of aganglionosis.

### ECM and cell adhesion components are altered in *Rdh10^trex/trex^* embryos

Our RNA-sequencing data also revealed that *Rdh10* loss-of-function results in altered focal cell adhesion and ECM composition (Figure 6). This is important as changes in adhesion and in the ECM influence cell migration and contribute to enteric nervous system defects as non-heritable factors in the etiology and pathogenesis of HSCR (72, 108–111). Many cell adhesion molecules expressed by vagal NCC promote their migration. For example, we found that *L1cam* is reduced in *Rdh10^trex/trex^* mutants, and interestingly, *L1cam* loss-of-function disrupts vagal NCC chain migration resulting in a delay in gut colonization (65). Furthermore, mutations in the *L1cam* gene were the first to associate a neural cell adhesion molecule with HSCR, (112) and its location on the X-chromosome is thought to contribute to male predominance in patients (113). Interestingly, *L1cam* acts as a modifier of the vagal NCC network through its interactions with *Sox10* and *Ednrb*, which together increase the incidence of aganglionosis (66, 86).(72, 108–111). Although it remains to be determined if RA directly regulates *L1cam*, our data provides further support for *Rdh10*-mediated RA signaling as a regulator of the genetic network that facilitates the migration of vagal NCC and their entry into the foregut.

The ECM is known to influence NCC migration and our RNA-sequencing data revealed considerable transcriptional changes in individual components in the interstitial and extracellular matrix. RA can regulate many of the affected matrix components including glycoproteins, laminin, fibronectin, tenascin-c, and collagen(68–70, 114–118). With respect to members of the collagen family we found that the levels of *Col1A1*, *Col1A2*, *Col2A1*, *Col3A1*, *Col4A2*, *Col4A6* and *Col6A3,* in addition to many others (Figure 8, Supplemental Table 3), were increased in *Rdh10^trex/trex^* embryos compared to controls. Increased COL I, COL III and COLVI have been observed surrounding the myenteric plexus in patients with HSCR(54) with high levels of COL IV in aganglionic gut segments (119). Additionally, the enrichment of collagen is consistent with the observation that an increase in ColVI restricts NCC migration in the pathogenesis of distal colonic aganglionosis(54). Furthermore COLVI has been posited to contribute to the high incidence of HSCR in patients with Down syndrome as several of its subunit genes are located on chromosome 21(54, 111).

The important role of collagen in HSCR pathogenesis can be explained in part through its abundance in the gut mesenchyme where it contributes to nearly 50% of ECM stiffness (120). Increased collagen concentration or deposition increases ECM stiffness which has subsequently been shown to restrict NCC migration in 3D gut explant cultures(72). Our data therefore reveals the importance of *Rdh10*-mediated RA signaling in regulating ECM composition, particularly with respect to collagen, to ensure proper vagal NCC migration. Thus, perturbation of cell adhesion and ECM composition in *Rdh10^trex/trex^* embryos may provide additional barriers to vagal NCC migration and entry into the foregut, during ENS formation and in the pathogenesis of intestinal aganglionosis.

## Conclusion

RDH10 oxidation of vitamin A plays a critical role in the synthesis of retinoic acid during embryogenesis(18, 19). RDH10 converts all-trans-retinol to all-trans-retinal (19, 23, 24) in a reversible manner with DHRS3, which serves as a key regulatory check-point in the pathway(19, 23, 24, 121–123). In this study, we uncovered the tissue specific role and mechanistic effects of Rdh10-mediated RA signaling during the early phases of ENS development. We showed that *Rdh10* is required during a narrow 48-hour window in which it regulates the microenvironment through which NCC migrate during their invasion of the foregut. This is fundamentally important for normal ENS formation, and when perturbed results in intestinal aganglionosis. Rdh10-mediated RA signaling functions in a paracrine manner to regulate the activity of key factors in the vagal NCC gene regulatory network including *Ret-Gdnf-Gfrα1*, together with *Phox2b*, *Pax3*, *Ednrb*, *Ecel1*, *Hoxb5* and *Col6a4*. Each of these genes has previously been associated with the etiology and pathogenesis of HSCR, through the critical roles they play in governing vagal NCC development. Rdh10-mediated RA signaling also influences cell adhesion and ECM composition with respect to laminin and collagen among many others, the perturbation of which may restrict NCC migration in association with the pathogenesis of HSCR. Collectively our data illustrates the pleiotropic effects of Rdh10-mediated RA signaling in regulating vagal NCC and their microenvironment during migration and entry into the foregut (Figure 9).

**Figure 9.**
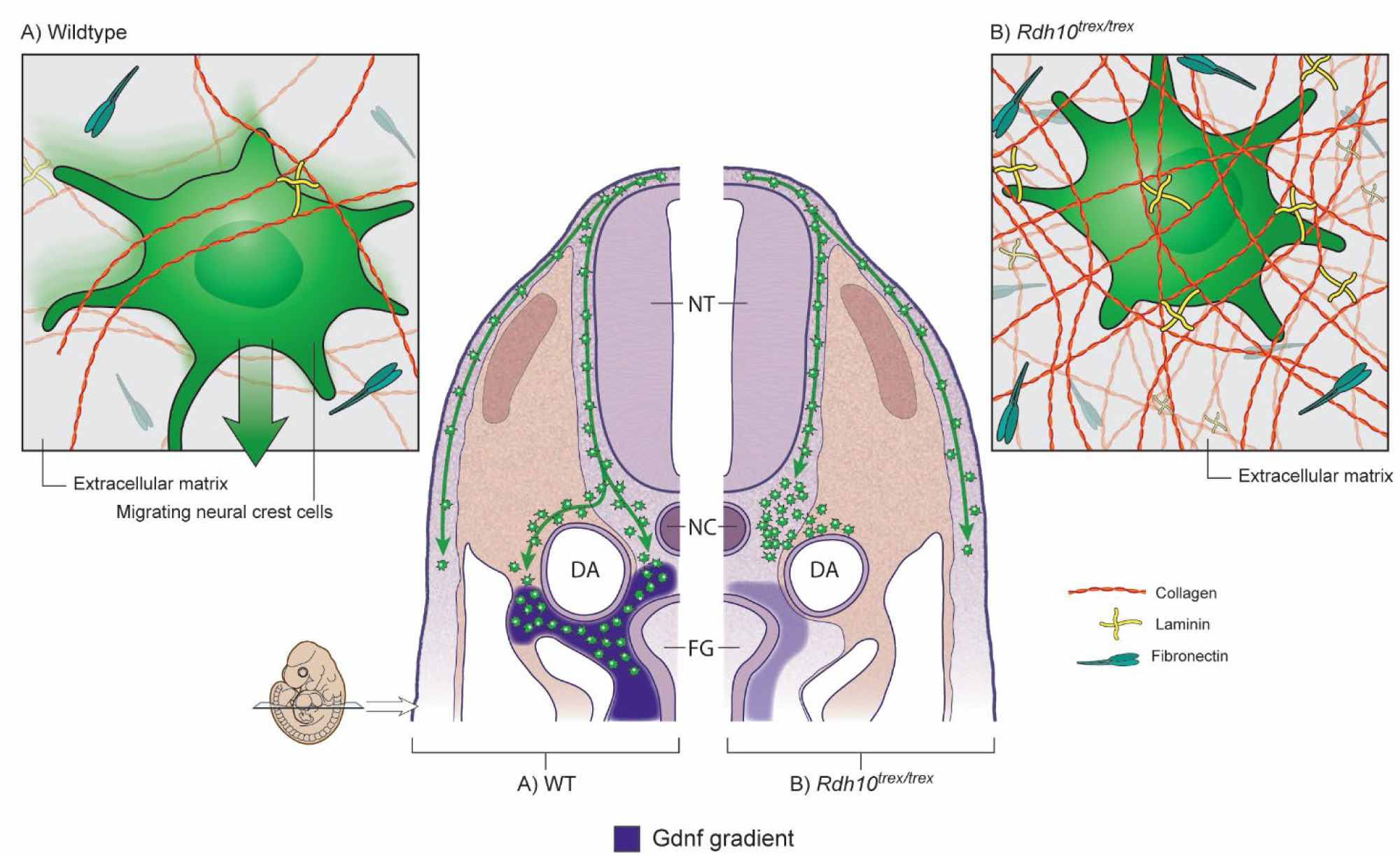
Proposed model of pathogenesis in *Rdh10^trex/trex^*. The point mutation in *Rdh10*reduces RA signaling, which restricts vagal neural crest cell migration into the foregut. Migration is disrupted by both a decrease in signaling from genes in an ENS signaling network, including *Ret*, *Gdnf* and *Gfra1*, and by an altered ECM composition, specifically increases collagen. Together, these changes limit access of migrating vagal neural crest cells into the foregut leading to incomplete gut colonization and total intestinal aganglionosis.

We investigated the spatiotemporal contributions of Rdh10-mediated RA signaling during ENS development and found that *Rdh10* is a novel key regulator of vagal NCC migration and invasion of the GI tract during ENS formation and in the pathogenesis of intestinal aganglionosis. Similarly, a recent study in mice showed that conditional activation of a dominant negative RAR in premigratory and migratory NCC, that is prior to their invasion of the foregut, also resulted in intestinal aganglionosis(83). Interestingly conditional activation at later stages, after foregut entry, resulted in only a mild hypoganglionosis phenotype. Furthermore, pharmacological and genetic modulation of the RA signaling pathway in zebrafish revealed that inhibition of RA signaling delays vagal NCC chain migration, resulting in distal aganglionosis(77). Conversely, exogenous RA expands the vagal NCC population leading to an increase in enteric neurons. Collectively, this illustrates the dynamic dual phase temporal contribution of RA signaling to vagal NCC development during ENS formation. RA signaling is firstly required for NCC invasion of the foregut, but then secondly is required after entry for the continued migration and differentiation of NCC throughout the GI tract. In addition, the RA signaling pathway also seems to be important for postnatal gut function as well, since patients with biallelic variants of *ALDH1A2*, present with defects in gut motility.(124) Our findings highlight the importance of *Rdh10*, vitamin A metabolism and RA signaling as both genetic and non-heritable environmental contributors to intestinal aganglionosis and possibly HSCR. Collectively, this demonstrates the importance of maintaining precise, appropriate levels of vitamin A and RA during pregnancy, and provides potential avenues for non-surgical prevention of HSCR by identifying and treating populations and pregnant mothers at high risk for vitamin A deficiency.

## Materials and Methods

### Mouse lines

Mice were housed in the Laboratory Animal Services Facility at the Stowers Institute for Medical Research and all animal experiments were performed in accordance with IACUC approved protocol 2022-143. *Rdh10^trex/+^*, *Rdh10*^βgeo/+^, *Aldh1a2^gri/+,^ RARE-lacZ, Mef2c-F10N-lacZ* and *ER^T2^Cre* (B6.129-*Gt(ROSA)26Sor^tm1(cre/ERT2)Tyj^*/J; Jax stock #008463) mouse lines were maintained as previously described (18–20, 125),(126). *Rdh10^flox/flox^*mice were created using Knockout Mouse Project Repository (KOMP) embryonic stem cells. Control embryos described in experiments using *Rdh10^trex/trex^* embryos were either wild-type or heterozygous littermates. Control embryos described in experiments using *Rdh10^fx/fx^* x Cre lines were either *Rdh10^fx/+^; ER^T2^Cre^+^*, *Rdh10^fx/+^* or *Rdh10^fx/fx^* littermates. For embryo staging, the morning of identification of the vaginal plug was defined as E0.5. All experiments were performed with a minimum N=5 unless specifically stated otherwise, and which were obtained from multiple litters for rigor and reproducibility. Although, sex was not specifically recorded for each embryo, our study included analyses of male and female embryos, with our cumulative findings reported.

### Gut organ culture

E11.5 embryos were dissected from the uterus and placed on ice cold PBS. Guts (esophagus, stomach, small and large intestines with accompanying mesentery) were isolated in fresh PBS, placed in L15 Leibowitz medium (Sigma) and then transferred onto a Millicell 0.4μm cell culture insert (Millipore) in a 6-well tissue culture dish (Millipore). The L15 medium was replaced with 400μL OptiMEM (Invitrogen) with 1mM L-Glutamine (Life Technologies Glutamax) and Penicillin/Streptomycin (Invitrogen). The guts were cultured in a 37°C incubator with 5% CO_2_ and the media was changed every 2 days as previously described. (127) For retinoid-supplemented cultures, guts were prepared as described above, and 1μM all-trans-retinoic acid (Sigma R2625) or 1μM all-trans-retinal (Sigma R2500) was added to the culture media (a 10^-6^ M concentration of RA produces morphological changes in 50% of cultured guts (128)).

### Organ Immunohistochemistry

E11.5-E18.5 embryos were dissected from the uterus into ice cold PBS. Isolated guts and cultured guts were fixed in 4% paraformaldehyde in PBS (PFA) overnight at 4^°^C. Guts were washed 3 x 5 minutes in PBS with 0.1% TritonX100 (PBT) at room temperature, followed by blocking and antibody hybridization in 10% heat-inactivated lamb serum in PBT. Non-specific antibody binding was blocked by a 20-minute incubation with blocking solution, after which primary antibody was applied overnight at 4^°^C in fresh blocking solution. Guts were then washed 5 x 1 hour in PBT at room temperature. Secondary antibody hybridization was performed in blocking solution overnight at 4^°^C, followed by 3 x 20-minute washes in PBT at room temperature. The guts were then stained with DAPI dilactate, 10 µg/ml in PBT, for 10 minutes or longer at room temperature and immersed into Vectashield with DAPI (Vector Laboratories) for storage and imaging.

### In situ hybridization

E8.5-E11.5 embryos were dissected from the uterus and into ice cold DEPC-PBS and then fixed in 4% paraformaldehyde (PFA) in phosphate buffered solution (PBS) overnight at 4°C. The following day, the embryos were dehydrated through an increasing methanol series diluted in DEPC-PBS 1% Tween (PBS-Tw), washing for 10 minutes at each step before being stored in 100% methanol at –20°C overnight. Embryos were rehydrated through a decreasing methanol series followed by 3 x 5 minute washes in PBS-Tw. The embryos were then treated with 10μg/mL Proteinase K in PBS-Tw without shaking. Proteinase K treatment time varied based on the age of the embryos. The embryos were washed 5 minutes in 2mg/mL glycine in PBS-Tw, then 2 x 5 minutes in PBS-Tw and were re-fixed in 4% paraformaldehyde/0.2% glutaraldehyde solution in PBS for 20 minutes. The embryos were then washed 3 x 5 minutes in PBS-Tw, rinsed in 65°C pre-warmed hybridization solution (for hybridization solution, MABT and NTMT recipes, see (129)), and placed in fresh pre-warmed hybridization solution for 2 hours at 65°C. Probe was added to fresh hybridization solution into which the embryos were incubated overnight at 65°C. The embryos were washed in pre-warmed hybridization solution 2 x 30 minutes, then 3 x 30 minutes at 65°C with Formamide Wash Solution (50% formamide, 1xSSC, pH4.5, 0.1% Tween). Next, the embryos were washed for 30 minutes in a 1:1 formamide wash solution: MABT, then 3 x 5 minutes MABT. The embryos were then blocked in MABT + 2% *Boehringer Blocking Reagent* (BBR) for 1 hour at room temperature, then an additional 1-2 hours in MABT + 2% BBR +20% heat-inactivated goat serum. The embryos were incubated overnight at 4°C in 1:2000 anti-DIG in MABT + 2% BBR + 20% heat-treated goat serum, rinsed, and then washed 2 x 15 minutes in MABT, then 5-7 x 1.5 hour in MABT, followed by an overnight MABT wash at room temperature. For anti-DIG antibody detection, the embryos were washed 3 x 10 minutes in NTMT, then incubated in the dark in NBT/BCIP reaction mix (3.375 µl/mL of 100mg/mL NBT and 3.5µL/mL of 50mg/mL BCIP per 1mL reaction mix) until the color reaction was deemed complete. Finally, the embryos were washed with PBS-Tw and fixed with 4% PFA/0.1% glutaraldehyde. *Gdnf in situ* hybridization required 50mg/ml Proteinase K in PBS-Tw at room temperature for 15 minutes, with occasional mixing. For color development, the embryos were incubated in the dark in NBT/BCIP reaction mix for up to 7 days with fresh reaction mix added every 24 hours.

### Whole embryo immunohistochemistry

E10.5-E11.5 embryos were dissected from the uterus and placed in ice cold PBS. Embryos were fixed in 4% PFA overnight at 4°C, then washed in PBS and transferred through an increasing methanol series into 100% methanol. Tissues and embryos to be immunostained were permeabilized in Dent’s bleach (MeOH:DMSO:30%H_2_O_2_, 4:1:1) for 2 hours at room temperature, then washed with 100% methanol and transferred through a decreasing methanol series into PBS. Blocking and antibody hybridization was performed in 0.1M Tris pH7.5/0.15M NaCl buffer. Non-specific antibody binding was blocked by a two-hour incubation in Tris/NaCl buffer with 3% Bovine Serum Albumin with gentle rocking. Primary antibody hybridization was performed in blocking buffer overnight at 4°C. Following primary antibody hybridization specimens were washed 5 x 1 hour in PBS at room temperature. Secondary antibody hybridization was performed in blocking buffer overnight at 4°C, followed by 3 × 20 minutes washes in PBS at room temperature. Specimens were stained with DAPI dilactate, 10 µg/ml in PBS, for 10 minutes or longer at room temperature or overnight at 4°C. Prior to imaging, immunostained specimens were cleared with Scale*A*2 for at least 5 days (130), or *Clear^T2^* for at least 2 days (131).Prior to imaging, immunostained specimens were cleared with Scale*A*2 for at least 5 days (130), or *Clear^T2^* for at least 2 days (131). Glass imaging wells were prepared by coating a Teflon O-ring with vacuum grease and affixing the ring to a glass depression slide. Scale*A*2 or *Clear^T2^* cleared specimens were placed into the well and the well filled with Scale*A*2 or *Clear^T2^*. A coverslip was then adhered to the vacuum grease coated O-ring.

### Antibodies

The following antibodies were used: βIII neuronal tubulin antibody (TUJ1), 1:1000, (MMS-435P or MRB-435P, Covance); GFP antibody, 1:500, (A6455 Invitrogen/Molecular Probes); Caspase 3 antibody, 1:500 (D175 Cell Signaling); AP2α antibody 1:500 (3B5, DSHB); Phox2b, 1:500 (gift from Dr Hideki Enomoto), ColVI 1:500 (Ab6588 Abcam); Sox10, 1:100 (AF2864 R&D System); Alexa Fluor 488, 1/500, (A21206 Invitrogen/Molecular Probes); and Alexa Fluor 546, 1/500, (A11030 Invitrogen/Molecular Probes).

### β-galactosidase staining

*Rdh10^trex/+^* mice were mated to *RARE-lacZ, Mef2c-F10N-lacZ* and *Wnt1;R26R* (126) mice and embryos/embryonic tissues were collected at E9.5-12.5. *Rdh10^βgeo/+^* mice were mated to FVB/NJ and embryos were collected at E8.5-10.5. Embryos were fixed with 2% Paraformaldehyde + 0.25% Glutaraldehyde, and then stained using a β-galactosidase staining solution kit (Chemicon/Millipore) according to the manufacturer’s instructions(132).

### Retinal gavage

312.5μg of all-trans-retinal (Sigma R2500) in 100µL corn oil (3.125mg/mL) was administered to pregnant females via oral gavage on the morning and night of E7.5, E8.5, and E9.5 (20).

### Tamoxifen gavage

5mg of tamoxifen (Sigma T5648) was administered to pregnant females via oral gavage at varying embryonic days post coitum in 100µL corn oil (50mg/mL). To prevent pregnancy loss, 1mg of progesterone (Sigma P3972) was co-administered (10mg/mL). To dissolve tamoxifen, 500mg of tamoxifen was initially dissolved in 1mL of ethanol and then 9mL of corn oil was added and warmed to 37°C until all the tamoxifen went into solution (approximately 8 hours). Using the *R26R; ER^T2^Cre* reporter line, 5mg tamoxifen + 1mg progesterone dosage results in complete Cre recombination within 24 hours of administration.

### Imaging

Confocal images were captured on an upright Zeiss LSM510 or LSM700 Pascal equipped with a 405 nm laser. For each specimen, a z-stack of images was collected and processed as a maximum intensity projection. Stacks of light microscope images were taken on a Leica MZ9.5 stereomicroscope and assembled in Helicon Focus.

### qRT-PCR

500ng of mouse RNA harvested from the vagal region of E9.0-E9.5 embryos delimited by the otic vesicle to the seventh somite was used as a template in a Superscript III (Life Technologies) reverse transcription kit reaction (40µl) utilizing random hexamer primers. The reaction volume was diluted 10 times and 2µl of cDNA was used in 10µl total qPCR reactions prepared with PerfeCTa SYBR Green FastMix, ROX (Quanta Biosciences) and cycled on an ABI 7900HT according to Quanta Biosciences standard protocols. Analysis of the fluorescence curves was performed using Life Technologies SDS2.4 software. All curves that showed errors as determined by the SDS2.4 software or that were above 35 Ct were discarded. The remaining Ct values were exported and analyzed using the Biogazelle qBase plus version 2.4. Three appropriate endogenous controls (M-value < .5) for each sample set were selected from a panel of five primer sets using geNorm in qBase plus. All primer sets (Supplemental Table 4) used were experimentally determined to be within 10% of ideal efficiency. Individual gene results were graphed with standard error bars, and relative expression (of all genes) results were graphed with 95% confidence intervals.

### RNA Sequencing

The vagal (somite 1-7) region of E9.5 embryos was used for RNA Seq (GSE166625) and QPCR experiments. Total RNA was extracted using a Qiagen RNEasy kit according to the manufacturer’s protocol. Samples were run on the Agilent 2100 BioAnalyzer and subsequently used to create polyA strand-specific libraries. Samples were sequenced on the Illumina HiSeq 2500 platform. Sequences for each RNA sample were aligned using TopHat (v2.0.10). The reference genome and annotation files used for the alignment were from the UCSC mm10 genome and the ensembl build 77. Expression levels for genes were calculated using R and Bioconductor. TopHat aligned reads were tabulated using the GenomicFeatures library, and differential expression was assessed using EdgeR(133). The experiment was performed in quadruplicate on 4 mutant and 4 wild-type littermates. Reads were counted for 55421 genes. However, after filtering for a sum of at least 10 reads across all samples, this left 20,883 genes remaining for analysis with edgeR. The top up regulated genes with ≥ 1.74 fold change and a p-value < 0.05 were examined for enrichment of GO terms from the Biological Process Ontology. Pathway enrichment analysis was also performed on all genes with an adjusted p-value of 0.05 or less (1992 genes) using Signaling Pathway Impact Analysis (SPIA), Database for Annotation, Visualization, and Integrated Discovery (DAVID)(134, 135), and ToppGene Suite(136, 137) developed at the Cincinnati Children’s Hospital Medical Center to assess pathway changes from databases that include the Kyoto Encyclopedia of Genes and Genomes (KEGG) Reactome, MSigDB C2 BIOCARTA, and Biosystems: Pathway Interaction Database. Ingenuity Pathway Analysis (IPA, Qiagen) software was also used to identify 128 genes that were differentially regulated that function in ECM organization, binding and/or adhesion. Additionally, select genes that were identified from the RNA Seq data that function in ENS formation, regulation of *Ret*, and HSCR pathogenesis were visualized in a gene interaction map using Cytoscape(138) software. The RNA Seq data referenced in this paper can be found in NCBI’s Gene Expression Omnibus(139, 140) with GEO Series accession number GSE166625 (https://www.ncbi.nlm.nih.gov/geo/query/acc.cgi?acc=GSE166625).

### Sirius Red staining

Embryos were collected at E9.5 and fixed with 4% PFA overnight at 4°C. The embryos were then rinsed in PBS and dehydrated in 70% ethanol and embedded in paraffin before being sectioned at 10µm. To identify collagen fibers in the anterior foregut, sections were stained with Sirius red and counter-stained with phosphomolybidic acid (PMA) to prevent auto-fluorescence during confocal imaging as previously described(141). The sections were imaged on an upright Zeiss LSM700 Pascal equipped with a 405 nm laser. 4 mutant and 5 wild-type embryos were analyzed across two litters. The fluorescent intensity of the Sirius red stain was measured across three consecutive sections of the anterior foregut for each embryo. Image J software(142) was used to quantify fluorescent intensity within a defined region of the anterior foregut. Statistical analysis between *Rdh10*^trex/trex^ and control littermates was performed using an unpaired student t-test with a calculated p-value of 0.0005.

## Acknowledgements

We thank members of the Trainor lab and the Neurogastroenterology community for their continual support and constructive feedback on this work. We greatly appreciate the Laboratory Animal Services Facility at the Stowers Institute for their care and maintenance of our mouse colonies. We thank the Molecular Biology core facility, particularly William McDowell for advice and analysis on qRT-PCR experiments. *RARE-lacZ* mice were a generous gift from J Rossant and we thank Dr. Hideki Enomoto for sharing the Phox2b antibody, Dr. Robert Heuckeroth for the Ret plasmid and Dr. Dipa Natarajan and Vassilis Pachnis for Ret, GDNF, Gfrα1 and EDNRB plasmids. We are extremely grateful to Mark Miller for generating the artwork in Figure 9. Research reported in this publication was supported by Stowers Institute for Medical Research (PAT), National Institute of Diabetes and Digestive and Kidney Diseases F30DK096842 (NEBT), and Madison and Lila Self Graduate Fellowship from the University of Kansas (SRS). Original data underlying this manuscript can be accessed from the Stowers Original Data Repository at http://www.stowers.org/research/publications/LIBPB-1603.

## Abbreviations

ENS: enteric nervous system
NCC: neural crest cells
HSCR: Hirschsprung disease
RA: retinoic acid
*Rdh10*: *retinol dehydrogenase 10*
ENU: N-ethyl-N-nitrosourea
RBP4: retinol binding protein 4
RAR: retinoic acid receptor
RXR: retinoid receptor
ECM: extracellular matrix composition
GO: gene ontology
SPIA: Signaling Pathway Impact Analysis
KEGG: Kyoto Encyclopedia of Genes and Genomes
IPA: Ingenuity Pathway Analysis
CBP: CREB-binding protein

