## Supplemental Figures for "Rdh10-mediated Retinoic Acid Signaling Regulates the Neural Crest Cell Microenvironment During ENS Formation"

**Supplemental Figure 1.** Vagal neural crest cell migration is restricted in *Rdh10*$^{trex/trex}$. *Sox10 in situ* hybridization labels migrating NCC in *Rdh10*$^{trex/trex}$ whole embryos at E9.5 (A,A',C,C') and E10.5 (B,B',D,D'). In control embryos at E9.5 (A,A') and E10.5b (B,B') vagal NCC staining is present in the neural tube and extends ventrally to the foregut region. *Rdh10*$^{trex/trex}$ embryos at E9.5 (C,C') and E10.5 (D,D') display staining evident in the neural tube without a vagal NCC stream. Black arrows notate foregut location.

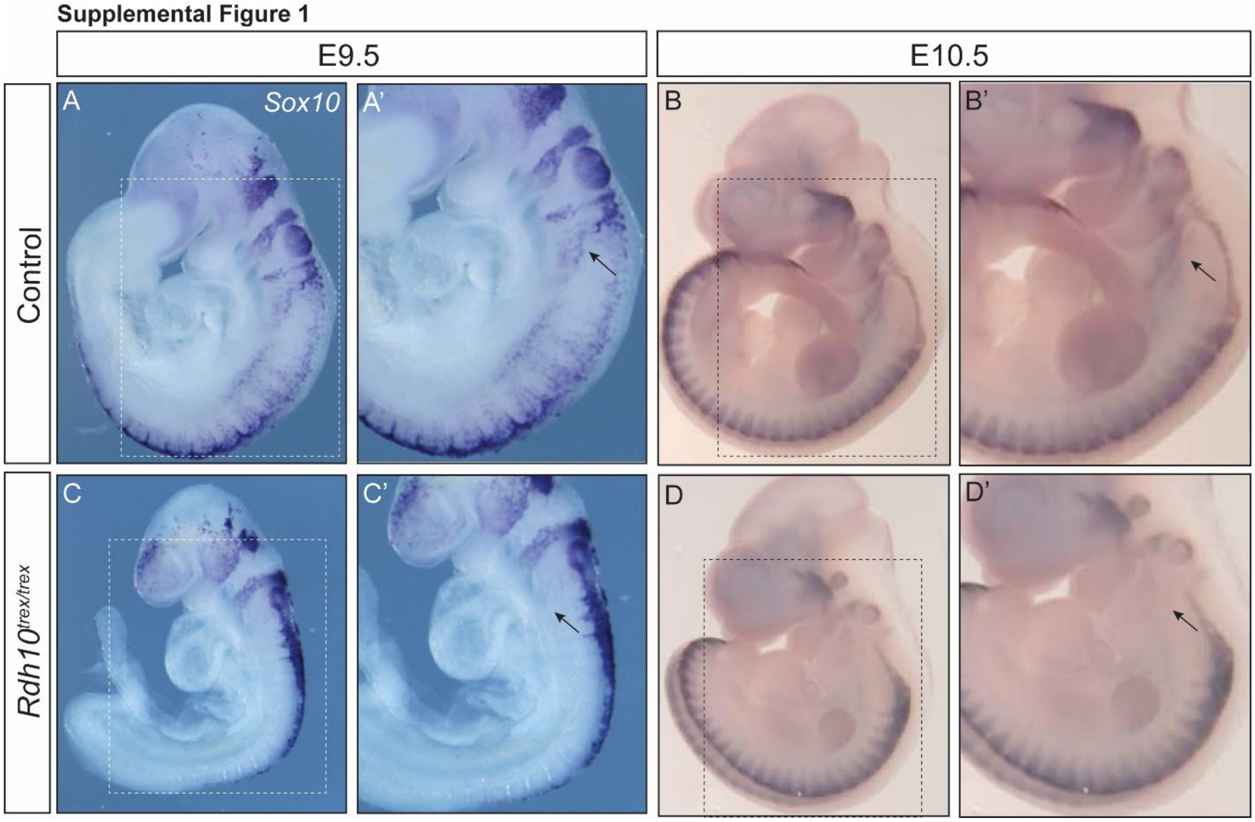

**Supplemental Figure 2.** Neural crest cells do have increased cell death staining during early colonization in *Rdh10*$^{trex/trex}$. NCC viability was assessed by immunostaining for Caspase-3 (red), and Ap2α (green) to label NCC. 10μM Cryosections of E9.5 in *Rdh10*$^{trex/trex}$ mutant and control embryos have reduced Ap2α+ neural crest cells near the foregut opening in mutant embryos (B,B') compared to littermates (A). Increased caspase 3+ cells is observed in mutant embryos (D,D') compared to control embryos (C,E), but caspase 3+ cells do not colocalize with Ap2α+ neural crest cells (F,F'). Rather, the cell death is restricted to the mesenchyme where NCC traverse. (nt) neural tube; (da) dorsal aorta; (fg) foregut.

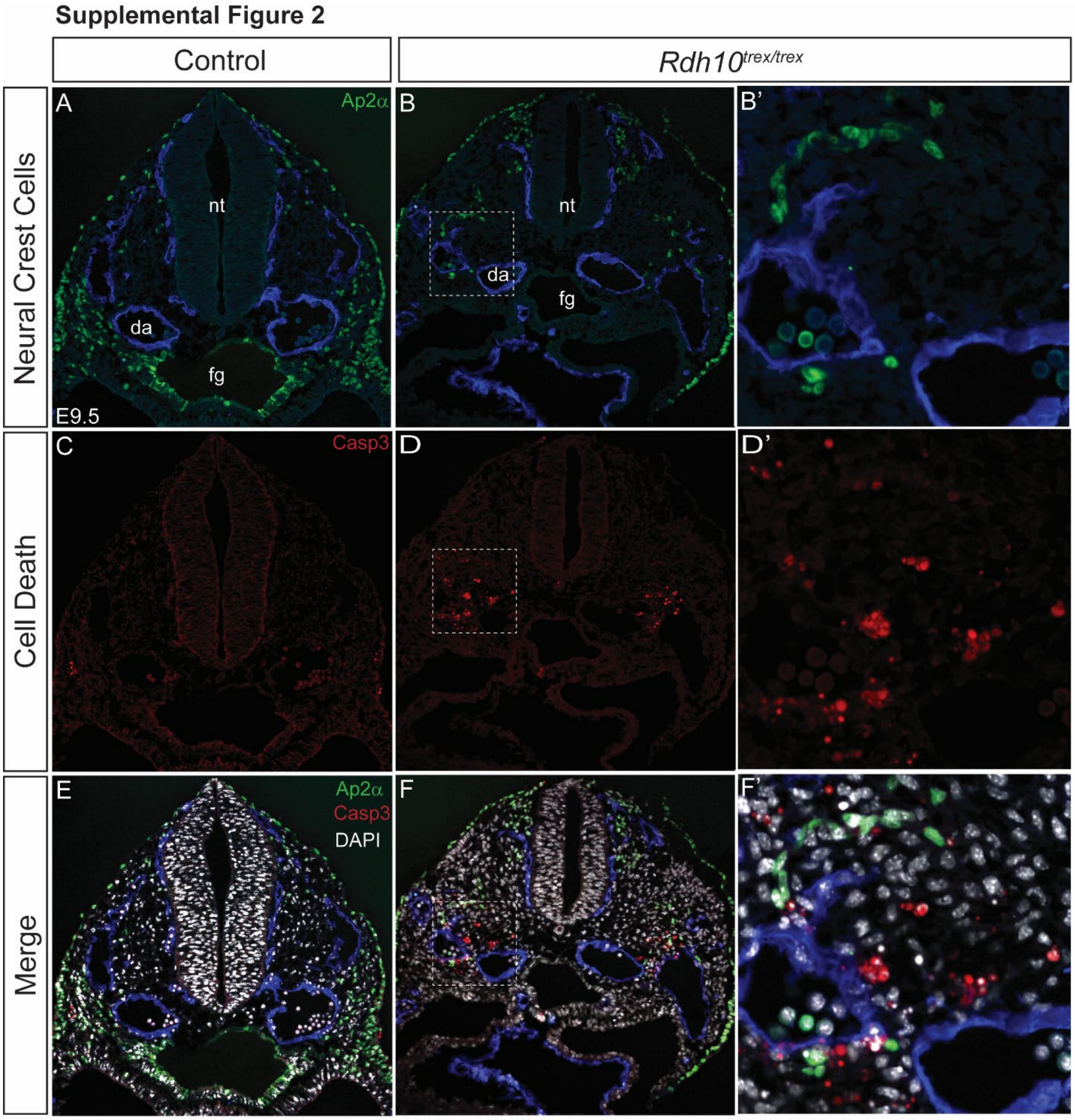

**Supplemental Figure 3**. RA signaling is required for vagal neural crest cell entry and colonization of the gut. (A-H) Whole guts dissected at E11.5 and cultured *ex vivo* in retinoid-free media 0 days (A,E), 2 days (B,F), 4 days (C,G) and 10 days (D,H). Cultured guts were immunostained with TUJ1 (red) and DAPI (blue) to assess enteric neuron differentiation. Control guts at 0 days (A) have mature neurons in the primitive gut from the stomach to the cecum. At 2 days (B), the control guts have neuronal differentiation through the length of the gut extending to the hindgut, and a dense, reticulated network throughout the full gastrointestinal tract at 4 days (C) and 10 days (d). $Rdh10^{trex/trex}$ guts at 0 days (E) have complete neuronal agenesis with little to no increase in colonization at 2 days (F), 4 days (G) or 10 days (H). (I-J') *In situ* hybridization of *Sox10* to label migrating NCC in the retinoid deficient $Aldh1a2^{gri/gri}$ mouse line at E9.5. Control embryos (I,I') have ventral NCC migration to the foregut, in contrast to mutant embryos (J,J') that lack vagal NCC migration. Black arrows notate expected foregut SOX10+ NCC stream. (s) stomach; (m) midgut; (h) hindgut; (c) cecum.

### Supplemental Figure 3

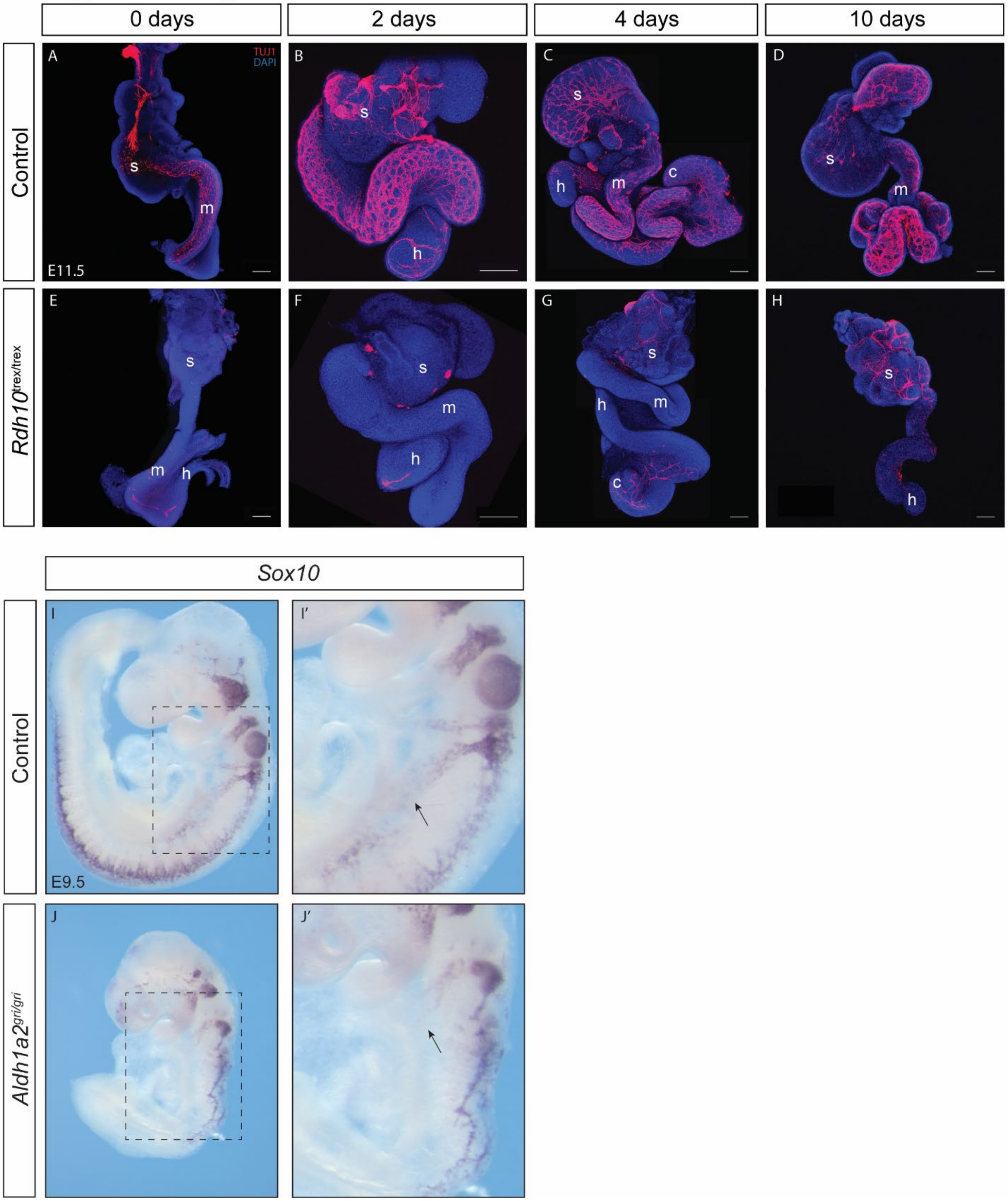

**Supplemental Figure 4.** *Rdh10*-mediated RA signaling is required prior to E11.5 for vagal neural crest cell migration. Whole guts dissected at E11.5 and cultured *ex vivo* for 4 days in either retinoid free media (A,D), or in the presence of all-trans-retinoic acid (B,E), or retinal (C,F). Cultured guts were immunostained with TUJ1 (red) and DAPI (blue) to assess enteric neuronal differentiation. Control guts (A) in retinoid free media display neuronal differentiation through each gut region leading to the hindgut. Guts cultured with all-trans-retinoic acid (B) display changes to the ENS that include punctate cells with few, short axonal projections. Treatment with retinal (C) caused changes producing fewer, disorganized axonal projections. *Rdh10*$^{trex/trex}$ guts in retinoid free media (D), treated with all-trans-retinoic acid (E), or retinal (F) were consistently aganglionic without noticeable improvement in neuronal differentiation.

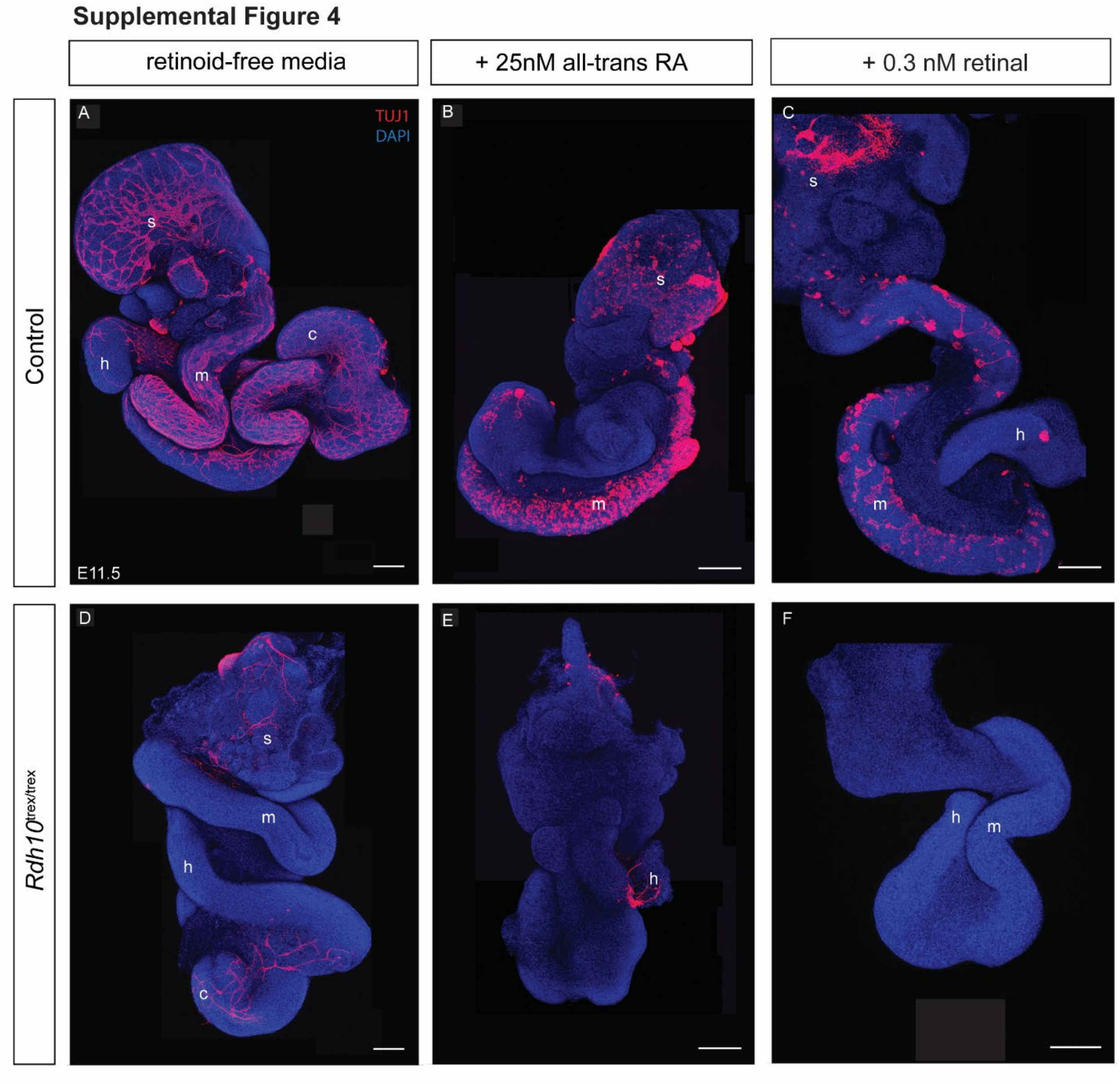

**Supplemental Figure 5.** *Rdh10* is not intrinsically required for vagal neural crest cell migration during gut colonization. A *Wnt1*-Cre mouse line was bred into the background of *Rdh10*$^{flox/flox}$ toconditionally delete *Rdh10* from NCC formed in the dorsal neuroepithelium. Whole embryos (A,B) and harvested guts (C-H) were immunostained for TUJ1 (red) and DAPI (blue) to assess neuronal differentiation. At E10.5, control *Rdh10*$^{fx/fx}$ (A) and mutant (B) *Rdh10*$^{fx/fx}$; *Wnt1*-Cre whole embryos both display neural formation present in the anterior foregut. At E15.5, mutant *Rdh10*$^{fx/fx}$; *Wnt1*-Cre guts (F,G,H) demonstrate grossly normal colonization with the same density and pattern of enteric neuron differentiation as the control *Rdh10*$^{fx/fx}$ guts in the duodenum, cecum and distal colon (C,D,E)

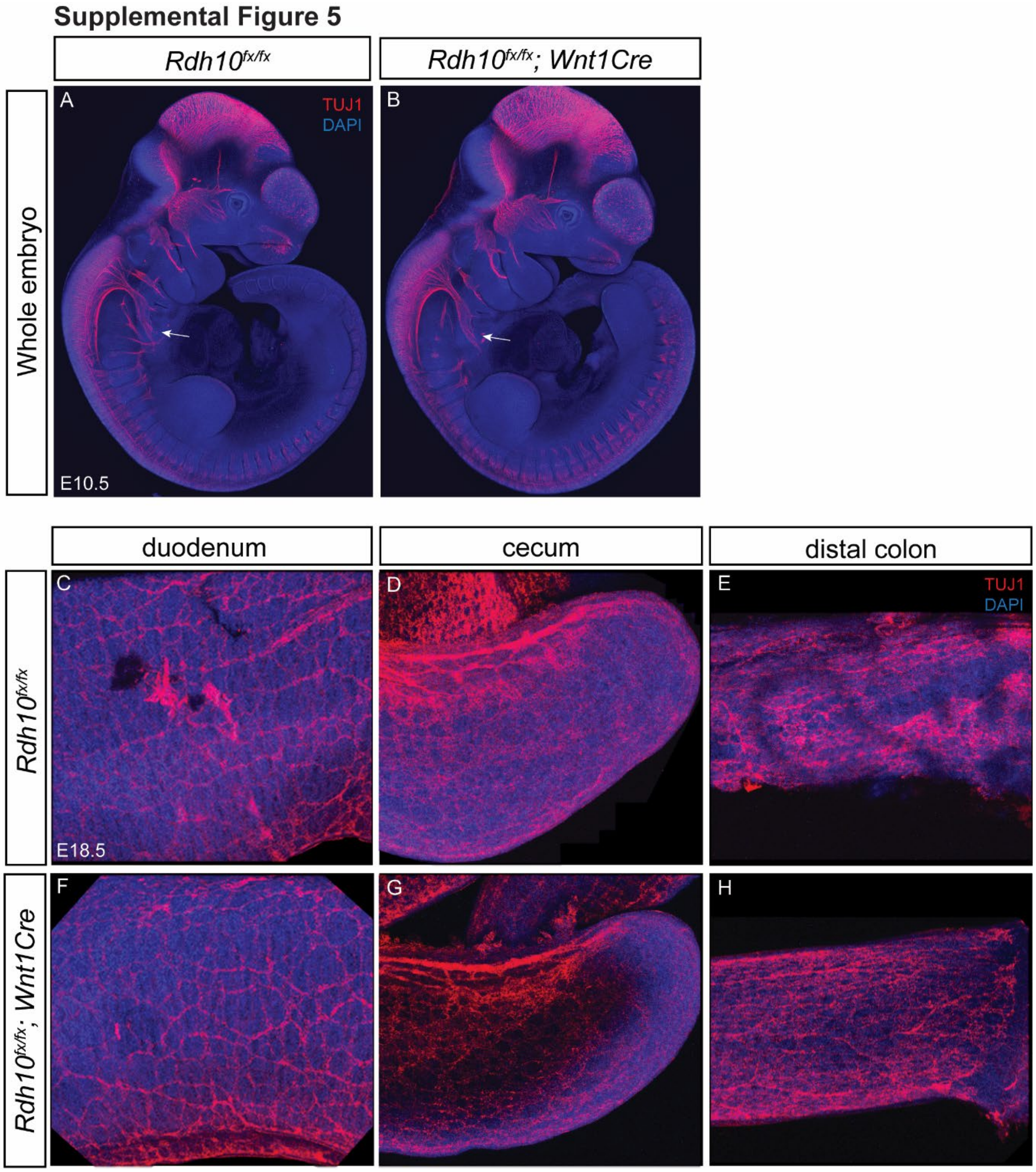

**Supplemental Figure 6.** *Gdnf activity disrupted in Aldh1a2$^{gri/gri}$* embryos. E9.5 control embryos display *Gdnf* expression within the foregut (Supplemental Figure 6 A,A', C, C'). In contrast, *Aldh1a2$^{gri/gri}$* embryos exhibit diminished *Gdnf* activity within the foregut region (Supplemental Figure 6 B,B', D, D').

Supplemental Figure 6

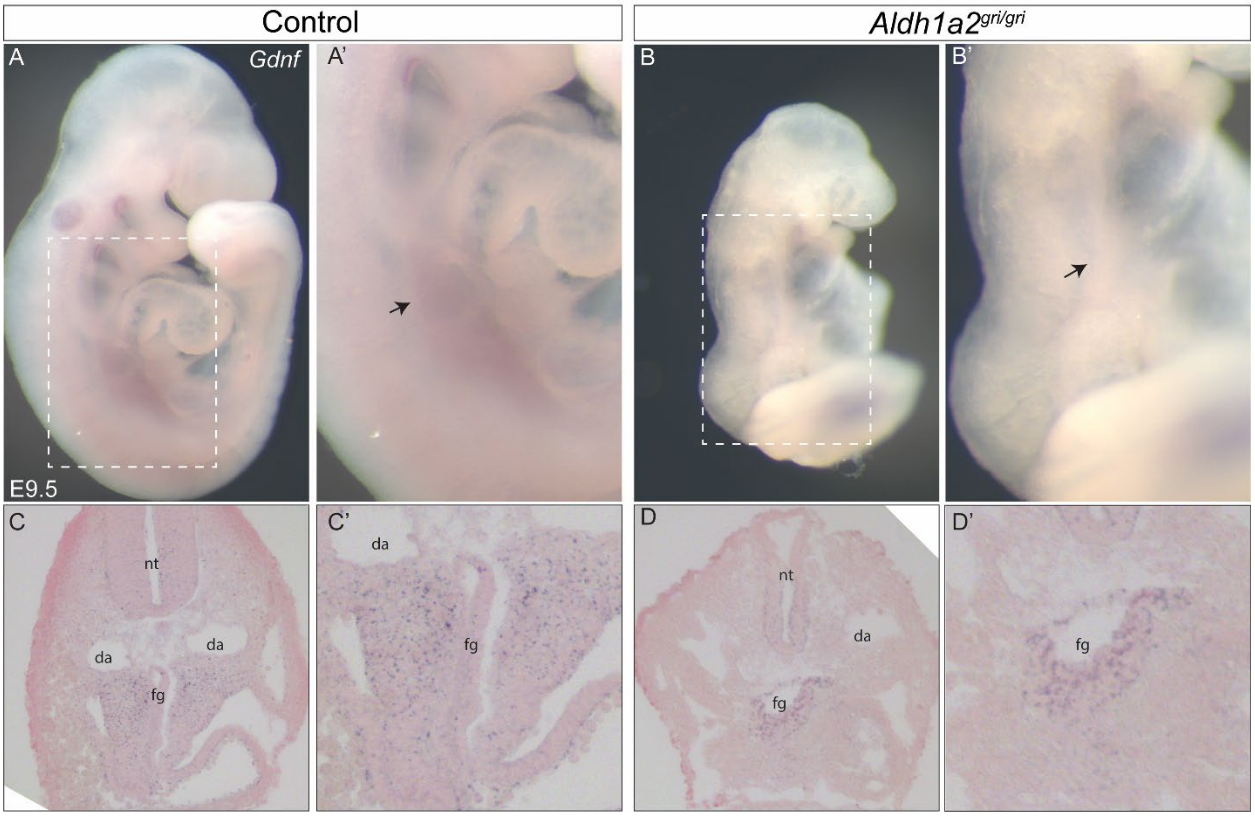

### Supplemental Tables

**Supplemental Table 1**. Genes differentially regulated in *Rdh10*$^{trex/trex}$ with established roles in ENS formation, *Ret* signaling, HSCR pathogenesis and HOX genes. LogCPM are the log counts per million.

| Gene | $\log_2$(trex/wt) | logCPM | p Value | $p^{adj}$ | Description |
|---|---|---|---|---|---|
| Col6a4 | 1.000 | -1.14 | 0.02964551 | 0.17674 | collagen, type VI, alpha 4 |
| Gfrα1 | -1.359 | 4.34 | 1.03E-09 | 1.54E-07 | glial cell line derived neurotrophic factor family receptor alpha 1 |
| Phox2b | -2.272 | 3.79 | 5.44E-08 | 4.49E-06 | paired-like homeobox 2b |
| Ecel1 | -2.279 | 1.29 | 3.89E-10 | 6.71E-08 | endothelin converting enzyme-like 1 |
| Nkx2.1 | -3.212 | 0.90 | 6.49E-17 | 5.90E-14 | NK2 homeobox 1 |
| Rarb | -1.173 | 5.76 | 2.65E-16 | 2.13E-13 | retinoic acid receptor, beta |
| Pax3 | -0.218 | 5.97 | 0.068515145 | 0.287395021 | paired box 3 |
| Hoxc9 | 6.064 | -0.57 | 2.03E-13 | 7.69E-11 | homeobox C9 |
| Hoxc8 | 5.742 | 3.06 | 4.93E-47 | 5.14E-43 | homeobox C8 |
| Hoxa9 | 2.742 | 2.45 | 4.84E-05 | 0.001374 | homeobox A9 |
| Hoxc6 | 2.560 | 3.50 | 1.39E-08 | 1.40E-06 | homeobox C6 |
| Hoxb9 | 2.422 | 6.99 | 7.94E-07 | 4.20E-05 | homeobox B9 |
| Hoxa7 | 2.365 | 3.54 | 0.00105391 | 0.016379 | homeobox A7 |
| Hoxb5os | 1.563 | 3.77 | 8.38E-07 | 4.39E-05 | homeobox B5 and homeobox B6, opposite strand |
| Hoxa6 | 1.312 | 0.36 | 0.00241179 | 0.030579 | homeobox A6 |
| Hoxb7 | 1.147 | 3.04 | 2.32E-06 | 0.000105 | homeobox B7 |
| Hoxd1 | 0.618 | 2.63 | 0.01884216 | 0.131731 | homeobox D1 |
| Hoxb8 | 0.432 | 5.35 | 0.00497795 | 0.051642 | homeobox B8 |
| Hoxc5 | -0.471 | 4.46 | 0.00422744 | 0.04598 | homeobox C5 |
| Hoxb3 | -0.611 | 6.70 | 4.02E-08 | 3.45E-06 | homeobox B3 |
| Hoxb5 | -0.710 | 4.41 | 5.27E-06 | 0.000212 | homeobox B5 |
| Hoxb1 | -0.769 | 4.02 | 5.53E-06 | 0.00022 | homeobox B1 |
| Hoxa1 | -0.860 | 4.63 | 2.27E-06 | 0.000103 | homeobox A1 |
| Hoxa4 | -0.985 | 4.55 | 7.02E-08 | 5.49E-06 | homeobox A4 |
| Hoxa10 | -3.451 | -0.93 | 0.0010054 | 0.015758 | homeobox A10 |

**Supplemental Table 2.** DAVID and ToppGene combined pathway enrichment analysis.

| Pathway | Database | p Value | Program |
|---|---|---|---|
| Extracellular matrix organization | BioSystems: Reactome | 2.96E-15 | ToppGene |
| Proteoglycans in cancer | KEGG | 2.56E-11 | DAVID |
| | MSigDB C2 BIOCARTA (v7.1) | 6.88E-11 | ToppGene |

| | | | |
|---|---|---|---|
| HIF-1-alpha transcription factor network | BioSystems: Pathway Interaction Database | 1.99E-10 | ToppGene |
| Focal adhesion | KEGG | 7.97E-11 | DAVID |
| | MSigDB C2 BIOCARTA (v7.1) | 3.82E-10 | ToppGene |
| ECM-receptor interaction | KEGG | 2.37E-10 | DAVID |
| | MSigDB C2 BIOCARTA (v7.1) | 1.09E-10 | ToppGene |
| Collagen biosynthesis and modifying enzymes | BioSystems: Reactome | 2.50E-08 | DAVID |
| Non-integrin membrane-ECM interactions | BioSystems: Reactome | 2.92E-08 | DAVID |

**Supplemental Table 3.** Genes differentially regulated in *Rdh10*$^{trex/trex}$ with functions in ECM organization, binding and adhesion identified by Ingenuity Pathway Analysis (IPA).

| **Symbol** | **Molecule Type** | **ID** | **Expr Log Ratio** | **Expr p-value** | **Expr False Discovery Rate (q-value)** |
|---|---|---|---|---|---|
| A2M | transporter | ENSMUSG00000030111 | 1.568 | 1.01E-03 | 1.58E-02 |
| Abi3bp | other | ENSMUSG00000035258 | 1.472 | 1.11E-07 | 8.04E-06 |
| ACER2 | enzyme | ENSMUSG00000038007 | 0.8 | 7.93E-06 | 2.98E-04 |
| ADAM12 | peptidase | ENSMUSG00000054555 | 0.373 | 3.39E-02 | 1.92E-01 |
| ADAM15 | peptidase | ENSMUSG00000028041 | 0.324 | 1.90E-02 | 1.32E-01 |
| ADAMTS12 | peptidase | ENSMUSG00000047497 | 0.764 | 1.66E-08 | 1.62E-06 |
| ADAMTS9 | peptidase | ENSMUSG00000030022 | 1.287 | 3.06E-17 | 2.90E-14 |
| ADAMTSL2 | other | ENSMUSG00000036040 | -0.888 | 1.91E-03 | 2.57E-02 |
| ATP7A | transporter | ENSMUSG00000033792 | 0.293 | 2.13E-02 | 1.43E-01 |
| BCAM | transmembrane receptor | ENSMUSG00000002980 | 0.299 | 1.86E-02 | 1.30E-01 |
| BCL3 | transcription regulator | ENSMUSG00000053175 | -0.955 | 3.68E-02 | 2.02E-01 |
| BGN | other | ENSMUSG00000031375 | 0.78 | 4.54E-04 | 8.40E-03 |
| BSG | transporter | ENSMUSG00000023175 | 0.758 | 1.19E-05 | 4.19E-04 |
| CAPN1 | peptidase | ENSMUSG00000024942 | 0.572 | 3.55E-03 | 4.06E-02 |
| CCDC80 | other | ENSMUSG00000022665 | 1.05 | 3.29E-09 | 4.07E-07 |
| CCN1 | other | ENSMUSG00000028195 | 0.501 | 6.18E-04 | 1.07E-02 |
| CCN2 | growth factor | ENSMUSG00000019997 | 0.814 | 7.38E-07 | 3.91E-05 |
| CD44 | other | ENSMUSG00000005087 | 0.856 | 1.26E-04 | 2.97E-03 |
| CD47 | transmembrane receptor | ENSMUSG00000055447 | 0.485 | 7.02E-03 | 6.65E-02 |
| CD82 | other | ENSMUSG00000027215 | 0.711 | 9.06E-03 | 8.01E-02 |

| | | | | | |
|---|---|---|---|---|---|
| CDH1 | other | ENSMUSG00000000303 | 0.452 | 1.54E-02 | 1.15E-01 |
| CDKN2A | transcription regulator | ENSMUSG00000044303 | 1.544 | 4.38E-02 | 2.22E-01 |
| CNTN2 | other | ENSMUSG00000053024 | -2.762 | 1.09E-10 | 2.09E-08 |
| COL18A1 | other | ENSMUSG00000001435 | 0.253 | 4.30E-02 | 2.19E-01 |
| COL1A1 | other | ENSMUSG00000001506 | 1.287 | 1.26E-07 | 8.94E-06 |
| COL1A2 | other | ENSMUSG00000029661 | 0.455 | 9.34E-04 | 1.49E-02 |
| COL2A1 | other | ENSMUSG00000022483 | 0.295 | 1.52E-03 | 2.17E-02 |
| COL3A1 | other | ENSMUSG00000026043 | 0.358 | 4.16E-02 | 2.15E-01 |
| COL4A1 | other | ENSMUSG00000031502 | 0.229 | 4.16E-02 | 2.15E-01 |
| COL4A2 | other | ENSMUSG00000031503 | 0.37 | 3.07E-03 | 3.65E-02 |
| COL4A5 | other | ENSMUSG00000031274 | 0.314 | 3.68E-02 | 2.02E-01 |
| COL4A6 | other | ENSMUSG00000031273 | 0.688 | 8.57E-05 | 2.15E-03 |
| COL5A1 | other | ENSMUSG00000026837 | 0.226 | 5.29E-02 | 2.48E-01 |
| COL5A2 | other | ENSMUSG00000026042 | 0.366 | 1.82E-03 | 2.50E-02 |
| COL6A3 | other | ENSMUSG00000048126 | 0.975 | 6.22E-05 | 1.67E-03 |
| Col6a4 | other | ENSMUSG00000032572 | 1 | 2.96E-02 | 1.77E-01 |
| COL8A1 | other | ENSMUSG00000068196 | 0.61 | 4.52E-02 | 2.26E-01 |
| COL9A2 | other | ENSMUSG00000028626 | 0.544 | 4.41E-02 | 2.22E-01 |
| COL9A3 | other | ENSMUSG00000027570 | 0.695 | 4.34E-02 | 2.21E-01 |
| CSGALNACT1 | enzyme | ENSMUSG00000036356 | 0.91 | 6.04E-05 | 1.63E-03 |
| CTSK | peptidase | ENSMUSG00000028111 | -0.831 | 2.92E-02 | 1.75E-01 |
| CTSV | peptidase | ENSMUSG00000021477 | 0.428 | 1.00E-02 | 8.54E-02 |
| CXCL12 | cytokine | ENSMUSG00000061353 | 0.236 | 3.23E-02 | 1.87E-01 |
| DCN | other | ENSMUSG00000019929 | 0.829 | 1.90E-02 | 1.32E-01 |
| DDR1 | kinase | ENSMUSG00000003534 | -0.264 | 6.65E-03 | 6.39E-02 |
| DDR2 | kinase | ENSMUSG00000026674 | -0.304 | 1.18E-02 | 9.53E-02 |
| DLL1 | enzyme | ENSMUSG00000014773 | -0.471 | 5.42E-03 | 5.51E-02 |
| Eda | other | ENSMUSG00000059327 | -0.521 | 1.29E-03 | 1.91E-02 |
| EGFL6 | other | ENSMUSG00000000402 | -0.765 | 4.57E-03 | 4.86E-02 |
| ELF3 | transcription regulator | ENSMUSG00000003051 | 0.754 | 1.08E-02 | 8.98E-02 |
| EMP2 | other | ENSMUSG00000022505 | 0.851 | 4.58E-03 | 4.87E-02 |
| ERO1A | enzyme | ENSMUSG00000021831 | 1.863 | 1.19E-07 | 8.49E-06 |
| EXOC8 | other | ENSMUSG00000074030 | -0.342 | 2.85E-02 | 1.73E-01 |
| F11R | other | ENSMUSG00000038235 | 0.46 | 2.63E-03 | 3.26E-02 |
| FBLN1 | other | ENSMUSG00000006369 | 0.365 | 3.60E-04 | 6.99E-03 |
| FBN1 | other | ENSMUSG00000027204 | 0.261 | 8.41E-03 | 7.62E-02 |
| FBN2 | other | ENSMUSG00000024598 | 0.322 | 2.65E-04 | 5.49E-03 |
| FGFR4 | kinase | ENSMUSG00000005320 | -0.655 | 1.51E-02 | 1.14E-01 |
| FN1 | enzyme | ENSMUSG00000026193 | 0.755 | 4.67E-09 | 5.57E-07 |
| FOXF2 | transcription regulator | ENSMUSG00000038402 | -0.637 | 8.45E-03 | 7.64E-02 |
| FREM1 | other | ENSMUSG00000059049 | 0.489 | 5.77E-06 | 2.28E-04 |
| GFOD2 | enzyme | ENSMUSG00000013150 | -0.291 | 3.96E-02 | 2.09E-01 |
| HAPLN1 | other | ENSMUSG00000021613 | 0.398 | 2.48E-03 | 3.12E-02 |

| | | | | | |
|---|---|---|---|---|---|
| HOXA7 | transcription regulator | ENSMUSG00000038236 | 2.365 | 1.05E-03 | 1.64E-02 |
| HSPG2 | enzyme | ENSMUSG00000028763 | 0.671 | 6.95E-08 | 5.46E-06 |
| HTRA1 | peptidase | ENSMUSG00000006205 | 0.649 | 1.23E-02 | 9.85E-02 |
| IBSP | other | ENSMUSG00000029306 | 2.456 | 7.00E-03 | 6.64E-02 |
| ICAM1 | transmembrane receptor | ENSMUSG00000037405 | 0.635 | 2.13E-03 | 2.79E-02 |
| IDUA | enzyme | ENSMUSG00000033540 | 1.031 | 3.60E-04 | 6.99E-03 |
| ITGA3 | other | ENSMUSG00000001507 | 0.336 | 1.05E-02 | 8.81E-02 |
| ITGA5 | transmembrane receptor | ENSMUSG00000000555 | 0.308 | 2.40E-03 | 3.05E-02 |
| ITGA6 | transmembrane receptor | ENSMUSG00000027111 | 0.331 | 1.39E-02 | 1.07E-01 |
| ITGA8 | other | ENSMUSG00000026768 | 0.336 | 4.80E-02 | 2.34E-01 |
| ITGAV | transmembrane receptor | ENSMUSG00000027087 | 0.25 | 2.42E-02 | 1.55E-01 |
| ITGB1 | transmembrane receptor | ENSMUSG00000025809 | 0.238 | 2.37E-02 | 1.52E-01 |
| ITGB4 | transmembrane receptor | ENSMUSG00000020758 | 0.75 | 1.39E-02 | 1.07E-01 |
| ITGB5 | other | ENSMUSG00000022817 | 0.422 | 6.72E-05 | 1.78E-03 |
| ITGB8 | other | ENSMUSG00000025321 | -1.448 | 3.26E-12 | 9.52E-10 |
| JAM2 | other | ENSMUSG00000053062 | -0.725 | 8.30E-05 | 2.10E-03 |
| KAZALD1 | other | ENSMUSG00000025213 | 0.939 | 1.49E-02 | 1.12E-01 |
| KDR | kinase | ENSMUSG00000062960 | 0.476 | 1.94E-04 | 4.26E-03 |
| L1CAM | other | ENSMUSG00000031391 | -0.677 | 2.10E-03 | 2.77E-02 |
| LAMA3 | other | ENSMUSG00000024421 | 1.067 | 1.25E-02 | 9.97E-02 |
| LAMA4 | enzyme | ENSMUSG00000019846 | 0.277 | 5.57E-02 | 2.55E-01 |
| LAMA5 | other | ENSMUSG00000015647 | 1.082 | 1.61E-09 | 2.21E-07 |
| LAMB1 | other | ENSMUSG00000002900 | 0.732 | 1.58E-05 | 5.29E-04 |
| LAMB2 | enzyme | ENSMUSG00000052911 | 0.341 | 3.81E-03 | 4.28E-02 |
| LAMB3 | transporter | ENSMUSG00000026639 | 1.278 | 9.36E-04 | 1.49E-02 |
| LAMC1 | other | ENSMUSG00000026478 | 0.474 | 5.74E-05 | 1.56E-03 |
| LAMC2 | other | ENSMUSG00000026479 | 0.595 | 1.81E-03 | 2.48E-02 |
| LAMC3 | other | ENSMUSG00000026840 | -1.795 | 1.49E-09 | 2.14E-07 |
| LGALS3 | other | ENSMUSG00000050335 | 1.808 | 1.18E-03 | 1.80E-02 |
| LOX | enzyme | ENSMUSG00000024529 | 0.806 | 4.03E-02 | 2.11E-01 |
| LRP1 | transmembrane receptor | ENSMUSG00000040249 | 0.568 | 2.87E-05 | 8.79E-04 |
| LYVE1 | transmembrane receptor | ENSMUSG00000030787 | 1.21 | 2.56E-03 | 3.19E-02 |
| MMP14 | peptidase | ENSMUSG00000000957 | 0.231 | 5.18E-02 | 2.44E-01 |
| MMP15 | peptidase | ENSMUSG00000031790 | -0.302 | 1.28E-02 | 1.01E-01 |
| MMP2 | peptidase | ENSMUSG00000031740 | 0.211 | 2.36E-02 | 1.52E-01 |
| NCAN | other | ENSMUSG00000002341 | -1.326 | 7.88E-08 | 6.03E-06 |
| NDNF | other | ENSMUSG00000049001 | 0.497 | 3.16E-04 | 6.33E-03 |

| | | | | | |
|---|---|---|---|---|---|
| NID2 | other | ENSMUSG00000021806 | 0.481 | 6.20E-04 | 1.07E-02 |
| NPHP3 | other | ENSMUSG00000032558 | 0.518 | 1.46E-03 | 2.11E-02 |
| OLFML2A | other | ENSMUSG00000046618 | 1.057 | 6.42E-05 | 1.72E-03 |
| PDGFRA | kinase | ENSMUSG00000029231 | 0.315 | 1.61E-03 | 2.27E-02 |
| PECAM1 | other | ENSMUSG00000020717 | 0.361 | 9.19E-03 | 8.09E-02 |
| POSTN | other | ENSMUSG00000027750 | 2.483 | 4.57E-10 | 7.64E-08 |
| PPFIA2 | phosphatase | ENSMUSG00000053825 | -2.291 | 3.66E-06 | 1.56E-04 |
| PXDN | enzyme | ENSMUSG00000020674 | 0.199 | 2.82E-02 | 1.71E-01 |
| RECK | other | ENSMUSG00000028476 | 0.463 | 5.15E-05 | 1.44E-03 |
| RXFP1 | G-protein coupled receptor | ENSMUSG00000034009 | 2.485 | 1.55E-07 | 1.07E-05 |
| SCUBE1 | transmembrane receptor | ENSMUSG00000016763 | -0.329 | 1.45E-03 | 2.10E-02 |
| SEMA3E | other | ENSMUSG00000063531 | 0.817 | 3.05E-03 | 3.64E-02 |
| SERPINE1 | other | ENSMUSG00000037411 | 1.299 | 3.34E-04 | 6.57E-03 |
| SIRPA | phosphatase | ENSMUSG00000037902 | 0.187 | 5.82E-02 | 2.61E-01 |
| SMOC2 | other | ENSMUSG00000023886 | 1.25 | 1.39E-06 | 6.72E-05 |
| SOX9 | transcription regulator | ENSMUSG00000000567 | 0.438 | 2.57E-05 | 8.01E-04 |
| SPARC | other | ENSMUSG00000018593 | 0.416 | 8.48E-04 | 1.38E-02 |
| SPINT1 | other | ENSMUSG00000027315 | 0.635 | 4.68E-03 | 4.94E-02 |
| SPOCK2 | other | ENSMUSG00000058297 | -0.812 | 1.10E-05 | 3.96E-04 |
| TEK | kinase | ENSMUSG00000006386 | 0.499 | 5.83E-04 | 1.02E-02 |
| TGFB2 | growth factor | ENSMUSG00000039239 | 0.509 | 1.86E-03 | 2.53E-02 |
| TGFBI | other | ENSMUSG00000035493 | 0.663 | 6.28E-08 | 5.02E-06 |
| THBS1 | other | ENSMUSG00000040152 | 0.382 | 2.30E-03 | 2.96E-02 |
| TIAM1 | other | ENSMUSG00000002489 | 0.341 | 1.10E-03 | 1.69E-02 |
| TNC | other | ENSMUSG00000028364 | 0.812 | 1.64E-05 | 5.47E-04 |
| TNFRSF11B | transmembrane receptor | ENSMUSG00000063727 | -0.853 | 2.33E-02 | 1.51E-01 |
| VTN | other | ENSMUSG00000017344 | -0.846 | 3.36E-02 | 1.92E-01 |
| WNT3A | cytokine | ENSMUSG00000009900 | 0.853 | 2.84E-04 | 5.79E-03 |

**Supplemental Table 4.** qRT-PCR primers.

| Target | Primer Sequence | Product size (bp) |
|---|---|---|
| GAPDH | F – TGGCCAAGGTCATCCATGA<br>R - CAGTCTTCTGGGTGGCAGTGA | 84 |
| GDNF | F – CTAAAGGAAAGGGTCAGGAG<br>R - TTGCTGCTCAGATGGATAG | 106 |
| RET | F – GCACAATTACAAGCTGATTCT<br>R - GAAATGGAGGACGAGGATAC | 130 |
| GFRα1 | F - GACTCCTGCAAGACAAATTAC<br>R - ACTGTGCCAATCAGTCC | 146 |

| | | |
|---|---|---|
| COL1A1 | F - CTAGACATGTTCAGCTTTGTG<br>R - GACTTCAGGGATGTCTTCTT | 93 |
| COL1A2 | F - CTGGACCAATGGGTTTAATG<br>R - AGCAGGTCCTTGGAAAC | 83 |
